# Organellar data sets confirm overall angiosperm relationships if problematic RNA-edit sites are accounted for in mitochondrial genomes

**DOI:** 10.1101/2025.06.09.658674

**Authors:** Wesley K. Gerelle, Matthias Jost, Isabel Marques, Don Les, Rodrigo Vallejos, Stefan Little, Brandon T. Sinn, Dmitry D. Sokoloff, Terry D. Macfarlane, Will Iles, Taylor Feild, Sarah Mathews, Michael Moore, Thomas L. P. Couvreur, Hervé Sauquet, Stefan Wanke, Sean W. Graham

## Abstract

**Premise:** Plastid-based data sets continue to play a major role in our understanding of early flowering-plant relationships, although organellar genomes of major lineages outside the monocots and eudicots remain under-sampled. A tendency of mitochondrial RNA-edit sites to mislead phylogenetic analysis in mixed transcriptomic/genomic data sets needs attention in angiosperm-wide studies, which only rarely consider mitochondrial genomes.

**Methods:** We compared mitochondrial- vs. plastid-based phylogenomic inferences, examined the effect of removing putative RNA-edit sites from mitochondrial data, and performed combined organellar analysis (plastid plus filtered mitochondrial genomes). We expanded taxon sampling for multiple angiosperm lineages for phylogenomic analysis using both organellar genomes, representing several poorly sampled lineages (in particular Degeneriaceae, Trimeniaceae) with smaller (few-gene) data sets.

**Results:** Plastid-based inferences recover well-supported relationships that align with and build upon previous studies, and recover well-supported internal relationships for two ANA-grade families (Hydatellaceae, Trimeniaceae) sampled for nearly all species. By contrast, unfiltered mitochondrial inferences of angiosperm phylogeny are generally poorly supported, and recover anomalous relationships compared to plastid-based inferences. However, removing putative mitochondrial RNA-edit sites dramatically reduces inter-organellar conflict and improves overall branch support.

**Conclusions:** We accounted for phylogenomic discordance between the two organellar genomes regarding overall angiosperm-wide relationships and filled in taxonomic gaps (poorly sampled lineages). Removing RNA edit sites substantially improves congruence in interorganellar inferences by effectively correcting a systematic bias in mitochondrial data. Uncertain relationships persist among five major mesangiosperm lineages in plastid-based inferences, but a clade comprising Chloranthales, Ceratophyllales and eudicots is well supported by filtered mitochondrial data.

## INTRODUCTION

Angiosperms (flowering plants) attract considerable attention from evolutionary and systematic biologists as they underpin human civilization and most terrestrial ecosystems (e.g., Benton et al., 2021 and references there), and are the most species-rich clade of plants (e.g., Christenhusz and Byng, 2016; Nic Lughadha et al., 2016) despite having a relatively recent origin and rise to dominance on the land (e.g., Friedman, 2009; Doyle, 2013; Magallón et al., 2013; Benton et al., 2021). The broad outline of angiosperm phylogeny has been well understood for several decades (e.g., Chase et al., 1993; Qiu et al., 1993, 1999; Mathews & Donoghue, 1999, 2000; Parkinson et al., 1999; Soltis et al., 1999, 2000, 2011; Barkman et al., 2000; Graham and Olmstead, 2000; Saarela et al., 2007), and was initially based on one to few-gene data sets, including the plastid genes *atp*B and *rbc*L, and the nuclear 18S rDNA gene. These legacy molecular results still underpin much of recent angiosperm classification (Angiosperm Phylogeny Group, APG, 1998, 2003, 2009, 2016), and thus of downstream phylogenetic analyses that rely on them, such as studies that integrate information from extinct and extant plants (e.g., Coiro et al., 2020). Despite our broad understanding of angiosperm phylogenetic relationships, additional refinements to our understanding of the broad sweep of their phylogenetic history have come from analyses based on whole plastid genomes (e.g., Jansen et al., 2007; Moore et al., 2010; Gitzendanner et al., 2018; Givnish et al., 2018; Li et al., 2019, 2021), transcriptomes (e.g. Wickett et al., 2014; One Thousand Plant Transcriptomes Initiative 2019), and other sources of genomic information, including studies based on target sequence data capture (e.g., Baker et al., 2021; Zuntini et al., 2024). However, it is clear that much work still remains to be done for undersampled lineages, and the ambiguous or uncertain parts of the angiosperm tree of life.

Numerous focused studies using phylogenetic and phylogenomic approaches have further refined our understanding of relationships in more recent angiosperm clades. For example, recent phylogenomic efforts have addressed the often rapidly evolving and difficult-to-place heterotrophic (holoparasitic and mycoheterotrophic) lineages in plant phylogeny, using evidence from transcriptomes and organellar genomes (e.g., Roquet et al., 2016; Lam et al., 2018; Jost et al., 2021; Lin et al., 2022; Timilsena et al., 2022; Chen et al. 2023). Despite these and similar advances in our knowledge of angiosperm phylogeny, major sets of relationships still resist satisfactory resolution, such as the precise relationships among five major mesangiosperm lineages (e.g., Fig. 1). These include uncertain relationships among the monogeneric order Ceratophyllales (*Ceratophyllum*), the monofamilial order Chloranthales (Chloranthaceae), and three very large angiosperm clades: the eudicots, magnoliids and monocots.

**Figure 1.**
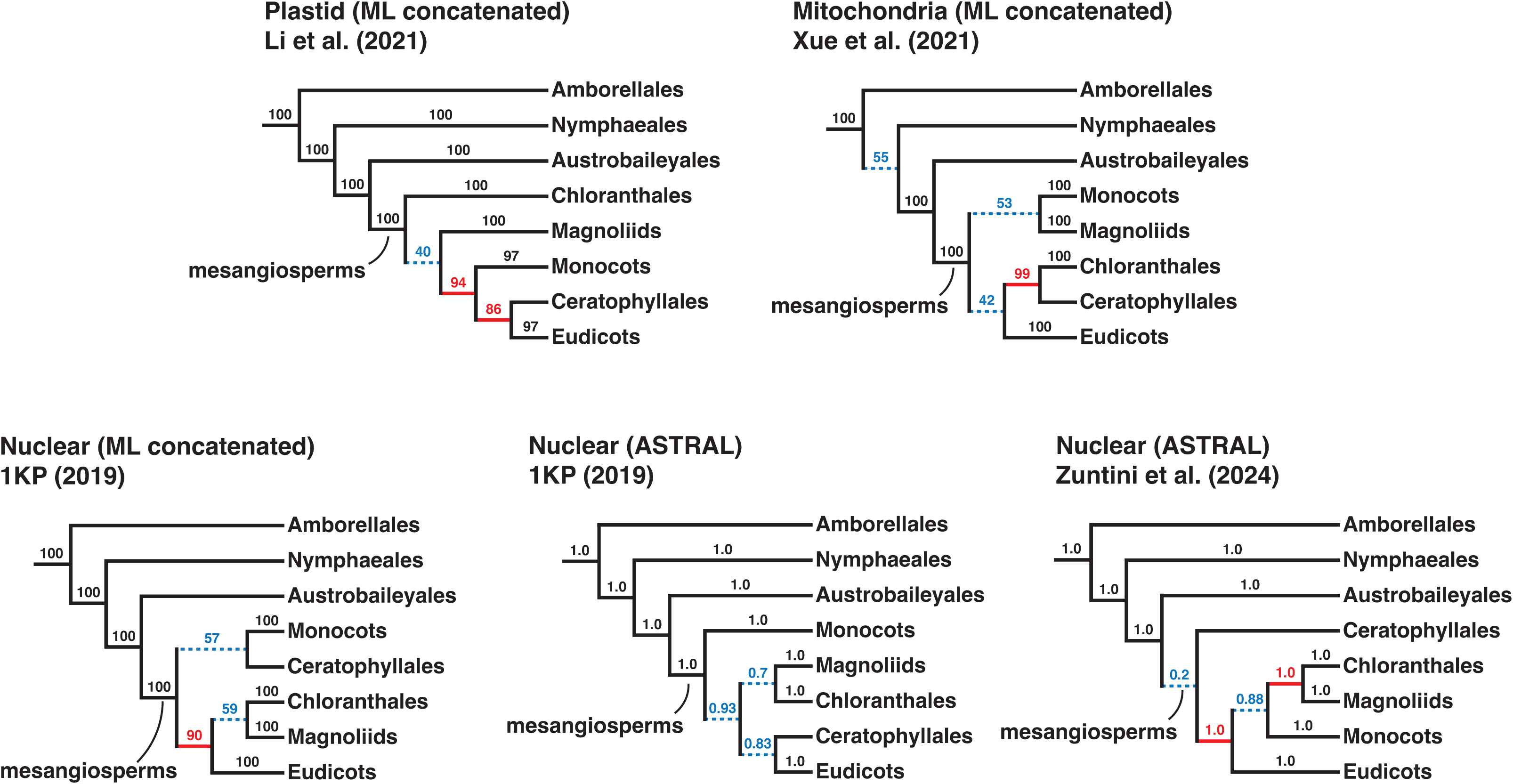
Schematic trees from recent studies showcasing conflicting topologies recovered in different studies; the study, phylogenetic approach (maximum likelihood analysis of concatenated data vs. ASTRAL coalescent-based analysis) and data source (plastid, mitochondrial or nuclear) are listed above each tree. Numbers above branches indicate support values (bootstrap or local posterior probability). Poorly supported internal branches (i.e., <70% bootstrap support or <0.95 posterior probability) are indicated with a dotted blue line, and blue font. Moderately to strongly supported clashes among the five trees (i.e., clashing branches both with at least 70% bootstrap support or 0.95 posterior probability somewhere in the five trees) are indicated with red lines, red font, and an asterisk.

Key lineages in major clades outside the monocots and eudicots also remain relatively under-represented in broad-scale phylogenomic analyses of angiosperm relationships (e.g., One Thousand Plant Transcriptomes Initiative, 2019; Dong et al., 2020; Li et al., 2021; but see Helmstetter et al., 2025). These include multiple lineages that comprise the four orders of the magnoliid clade (Canellales, Laurales, Magnoliales and Piperales), and individual families in these and other orders. Some taxa that are highly relevant to understanding early angiosperm evolution have hardly been examined using molecular data (e.g., Trimeniaceae, Austrobaileyales), or remain poorly represented in phylogenomic data sets (e.g., Hydatellaceae, Nymphaeales). For those that have been examined in more detail, uncertainty persists about major sets of relationships within orders. For example, within Laurales the exact relationships among Monimiaceae, Hernandiaceae and Lauraceae have been controversial, and all three possible relationships among these families have been found in other studies based on multiple lines of data, generally with weak to moderate bootstrap support (Qiu et al., 1999, 2000, 2005, 2006, 2010; Renner, 1999; Renner and Chanderbali, 2000; Savolainen et al., 2000; Hilu et al., 2003; Zanis et al., 2003; Soltis et al., 2011; Massoni et al., 2014—but see Doyle and Endress, 2000, 2010, and Sauquet et al., 2003). In Magnoliales, the precise positions of Degeneriaceae and Himantandraceae are also unclear, and multiple poorly supported relationships have been reported concerning these taxa and the other Magnoliales families (i.e., Annonaceae, Eupomatiaceae, Magnoliaceae and Myristicaceae; see Massoni et al., 2014, and Helmstetter et al., 2025).

While plastid data sets have provided the broad framework for modern angiosperm classification, mitochondrial-based phylogenetic studies have been used much less frequently, although Qiu et al. (1999, 2000, 2006, 2010) used small sets of mitochondrial genes to infer early angiosperm relationship, and Xue et al. (2022) considered whole mitochondrial genomes for this purpose in a recent broad-scale study. An apparent hesitancy to use this genome may reflect the concern that mitochondrial genes have an extremely slow substitutional rate in general (e.g., Wolfe et al., 1987), with the result that individual genes generally provide little phylogenetic information. Nonetheless, genome-scale mitochondrial data sets concatenating all or most of the protein-coding genes have recently permitted robust phylogenetic inference of higher-order relationships (e.g., Bell et al., 2020; Soto Gomez et al., 2020; Dong et al., 2022; Klimpert et al., 2022; Lin et al., 2022; Xue et al., 2022), supporting their use in higher-order phylogenomic inference.

A major concern in using mitochondrial genome data is that transcripts from the land-plant mitogenome can have high levels of RNA editing (C to U, or U to C) compared to plastid transcripts (e.g., Knoop, 2023; Takenaka et al., 2023). These sites may present a significant challenge for phylogenetic reconstruction, if not properly accounted for when combining sequences derived from transcriptomes (RNAseq) vs. genomic DNA (e.g., genome-skim or target-sequence capture for the latter). Mixing transcribed and genomic data sources may lead to clustering of lineages based on the respective nucleic-acid type (RNA vs. DNA), rather than by phylogenetic history (Bowe and DePamphilis, 1996). This effect occurs when these two types of data are included together in phylogenetic inference without correcting for putative edit sites. In this situation, RNA-based sequences in this situation may mistakenly act as false evidence of ancestry (i.e., false synapomorphies), artefactually pulling them sequences together in phylogenetic inference, in opposition to taxa represented by genomic sequences (e.g., Bell et al., 2020; Dong et al., 2022). This type of systematic error was first described by Bowe and DePamphilis (1996) for two mitochondrial genes, who cautioned that DNA and RNA-derived data should not be mixed in phylogenetic studies (e.g., as in One Thousand Plant Transcriptomes Initiative, 2019, for example). Fortunately, alignment sites prone to RNA-editing tend to be conserved evolutionarily (e.g., Lenz and Knoop, 2013) and can be predicted for taxa where genomes and transcriptomes are both available for the same species (i.e., sites unedited in genomic data vs. edited in transcriptome data). Qiu et al. (2006, 2010) also compared phylogenetic analyses with RNA edits sites included or excluded and found little effect; however, they did not mix sequences derived from RNA and DNA in their analyses, focusing only on the latter.

Here we expand the density of major angiosperm lineages sampled for organellar genomic data, using a variety of published sources and new data. Bell et al. (2020) recently demonstrated that simply removing affected alignment columns in data sets with transcriptomes and genome sequences can be an effective strategy to correct for RNA edits in phylogenetic inference of combined transcriptome and genome data, even when only a subset of taxa can be assessed to predict shared RNA edit sites across taxa. This is the approach we use here: we infer putative RNA edit sites by comparing genomic and transcriptomic data for a core set of taxa, and remove the corresponding sites across the aligned taxa. A similar principle is used in the software package PREPACT (Lenz and Knoop, 2013). We then perform phylogenomic inference for both organellar genomes, individually and combined, after filtering out sites that are likely to be RNA edits in the mitochondrial data.

In addition, in cases where genome-scale data are unavailable for several key lineages, we included lineages represented by small (few-gene) plastid data sets. This facilitates species-level sampling in particular for two families (Hydatellaceae, Trimeniaceae) in the ANA grade in particular, and for addressing the placement of Degeneriaceae within Magnoliales. The overall aims of our study are: (1) to use organellar genome-scale data (plastid and mitochondrial) to re-visit phylogenetic inference of the early angiosperm radiation, including the relative branching order of major lineages defining the earliest phylogenetic splits in the flowering plants, along with improved taxon sampling in multiple ANA-grade and magnoliid lineages; (2) to identify and account for the possible impact of RNA editing on phylogenomic inference when using mixed genomic/transcriptomic sources of mitochondrial data, and; (3) to use combined analyses of plastid and mitochondrial genome data to infer overall angiosperm relationships.

## MATERIALS AND METHODS

### Taxon sampling, data recovery and sequencing

We assembled plastid and mitochondrial gene sets for taxon sets that focused on the major lineages of angiosperms: Amborellales, Nymphaeales and Austrobaileyales (collectively known as the ANA grade), magnoliids, Chloranthales, and Ceratophyllales, in addition to a lighter sampling of representative orders from eudicots and monocots, and a collection of gymnosperm outgroups (Table S1). The angiosperms are represented here by 60 and 80 newly sequenced taxa (see below) for the mitochondrial and plastid data sets, respectively. We generated new data for both data sets using shotgun sequencing (e.g., Cronn et al., 2008; Moore et al., 2010; Ross et al., 2016). We added these to sequence data from 53 and 146 additional angiosperm taxa (for mitochondrial and plastid data sets, respectively). We retrieved data for additional taxa from published data sets from GenBank, 1KP (One Thousand Plant Transcriptomes Initiative, 2019), Rossetto et al. (2015), Hoekstra et al. (2017), Li et al. (2019, 2021), and other sources, including floral transcriptomes produced using methods described in Couvreur et al. (2019); the sources are summarized in Table S1. We extracted plastid and mitochondrial genes from both transcriptomes and genomes for individual taxa by applying a custom set of Python scripts (github.com/wesleykg/pull_out_genes) and BLASTN and tBLASTX (Altschul et al., 1990). For the plastid dataset, we used plastid sequences from the following GenBank samples as query sequences: *Amborella trichopoda, Calycanthus floridus, Ceratophyllum demersum, Chloranthus spicatus*, *Drimys granadensis, Illicium parviflorum, Liriodendron tulipifera, Nuphar advena* and *Piper cenocladum* (see Table S1 for source information). For the mitochondrial data set we used *Liriodendron tulipifera* (from GenBank; see Table S1), and *Asarum minus* and *Saruma henryi* (for the latter two assembled here from the same specimens used to produce the GenBank plastid data shown in Table S1, NC_037503 and NC_039933, respectively).

We represented members of a few key lineages that are not represented by plastid or mitochondrial genomes with small few-gene sets taken from GenBank (i.e., *Degeneria*, *Galbulimima*; Table S1) or newly sequenced here using Sanger-based approaches and primer sets (Graham and Olmstead, 2000, and see Iles et al., 2012), in particular for multiple species in Hydatellaceae and Trimeniaceae (see Table S1). However, in the latter cases, at least two species in both Hydatellaceae and Trimeniaceae are represented by full plastid genomes (Table S1). For the mitochondrial dataset, we included two *Degeneria* species with only one or two genes, respectively. In total, the few-gene data sets in both the plastid and mitochondrial dataset range from one gene (*Degeneria roseiflora*) to eight genes total (i.e., for some species of *Trimenia*, see Table S1, Table S6; we used the approach in Vaidya et al., 2011, to prepare the latter table).

The plastid sampling includes all families and genera in the ANA grade and Chloranthales, nearly all families, and 97 genera of magnoliids. Fewer taxa were available from published sources for the mitochondrial genome. However, our overall sampling for this genome includes a taxonomically broad subsampling of focal lineages represented in both organellar genomes (i.e., almost all genera in the ANA grade, two of four genera of Chloranthales, and nearly all families; it includes 63 genera in the magnoliids). We excluded the holoparasitic lineage Hydnoraceae as it is a very rapidly evolving lineage whose placement is now quite well understood (see Jost et al., 2021). We follow the family- and order-level classification in APG (2016), except that we recognize Asaraceae and Lactoridaceae in Piperales, following Jost et al. (2021).

### Data assembly, filtering and alignment

The final mitochondrial and plastid datasets include 141 and 275 taxa, with 26 and 47 gymnosperm outgroups, respectively; they comprise 42 and 77 protein coding genes, respectively (Table S6). Our assembly methods broadly follow Moore et al. (2010) and Ross et al. (2016). In particular, we assembled raw reads for newly sequenced genomes using the *de novo* assembly tool in CLC Genomics Workbench 6.0 and 11.0 (CLC bio, Aarhus, Denmark) allowing for automatic word and bubble size, the auto-detection of paired distances, setting the minimum contig length to 150 bp, and also created read mappings of raw reads back to the contigs, setting the length and similarity fraction to 0.85. Some of the assemblies from 1KP pieced together either barely or not fully as overlapping scaffolds with each other, but otherwise spanned the full length of an ORF (open reading frame). These include several cases where the overlap was one or a few nucleotides long (also the case in Wickett et al., 2014, and the One Thousand Plant Transcriptomes Initiative, 2019). In a few cases where the overlapping region contained divergent bases, we excluded the transcript with lower sequencing depth, retaining the other for further analysis. Partial genes from non-overlapping BLAST hits were merged into one sequence with gaps included across the unretrieved portions.

We aligned individual plastid or mitochondrial genes using MAFFT v7 (Katoh et al., 2002; Katoh & Stanley, 2013) using the L-INS-i algorithm (“localpair” option) and a “maxiterate” value of 1000. We manually refined sequences in AliView (Larsson, 2014) following Graham et al. (2000), and concatenated the single gene alignments using a custom Python script (Nathaniel Klimpert, University of British Columbia; https://github.com/nklimpert/Phylogenetics_Scripts). The final alignments are on FigShare, see Data Availability Statement. We ran preliminary phylogenetic analyses using RAxML version 8.2.12 (Stamatakis, 2014) on individual genes (GTR+G model with 10 best tree searches and 100 rapid bootstraps) to visually identify and remove problematic sequences that represent contaminants (these typically had extremely long branches; data not shown); for example, for genes recovered from the 1KP data, we identified several prokaryotic and animal (insect) scaffolds, both plastid and mitochondrial for ribosomal DNA (rDNA) genes, and *cob* and *cox1* (from the mitochondrial genome). We retained relatively rapidly evolving sequences for several lineages with recognized long branches, including mitochondrial sequences for *Acorus* (Qiu et al., 2006; Hertweck et al., 2015) and plastid sequences for Poales and Gnetales (Duvall et al., 1993; Doyle, 1998).

### Accounting for putative RNA edit sites in phylogenetic inferences of mixed genome/transcriptome data

We identified possible RNA-edited sites (C-to-U or U-to-C edits) in mitochondrial and plastid data by comparing paired genomic and transcriptomic data for sequences from *Amborella trichopoda* and also species representing three of the four magnoliid orders: *Eupomatia bennettii, Gyrocarpus americana, Houttuynia cordata, Magnolia maudiae, Myristica fragrans, Saruma henryi* and *Saururus cernuus* for the mitochondrial dataset; and the same set of species plus *Annona muricata, Calycanthus floridus* in the magnoliids, and *Austrobaileya scandens, Ceratophyllum demersum, Nuphar advena,* and *Trochodendron aralioides* from other major lineages for the plastid dataset (see Table S1 and footnotes). This approach identifies putative RNA edit sites, as the genomic sequences are unedited and transcriptomic sequences include RNA edits. Similar approaches like PREPACT (Lenz and Knoop, 2013) consider additional information to identify probable edits (e.g., whether reverting the edit site in the transcriptomic DNA restores an amino acid residue coded by the genomic DNA). Here we used only direct comparisons to predict RNA edit sites, a simple approach that proved effective in correcting biases due to RNA editing in land-plant data sets (Bell et al., 2020). We used the resulting putative RNA edit sites (C-to-U or U-to-C) to create versions of the main plastid and mitochondrial matrices in which all nucleotide positions in an alignment column were converted to an “N” (= unknown nucleotide) if there was at least one corresponding DNA-to-RNA (genome to transcriptome) U-to-C or C-to-U difference for at least one reference taxon. We recorded the length of overlap between transcriptomic vs. genomic sequences for each individual gene in a given species, and the number of disagreements between transcriptomes and genomes in their overlapping sequences (for all categories of nucleotide difference, used to make the inferences summarized in Fig. 2). We also assessed how many of the putative edit sites are shared with the other species (Tables S3, S9), and for the overlapping mitochondrial regions for individual species (Table S4). We made these comparisons using a custom R script (https://github.com/wesleykg/ANAM_RNA_editing/).

**Figure 2.**
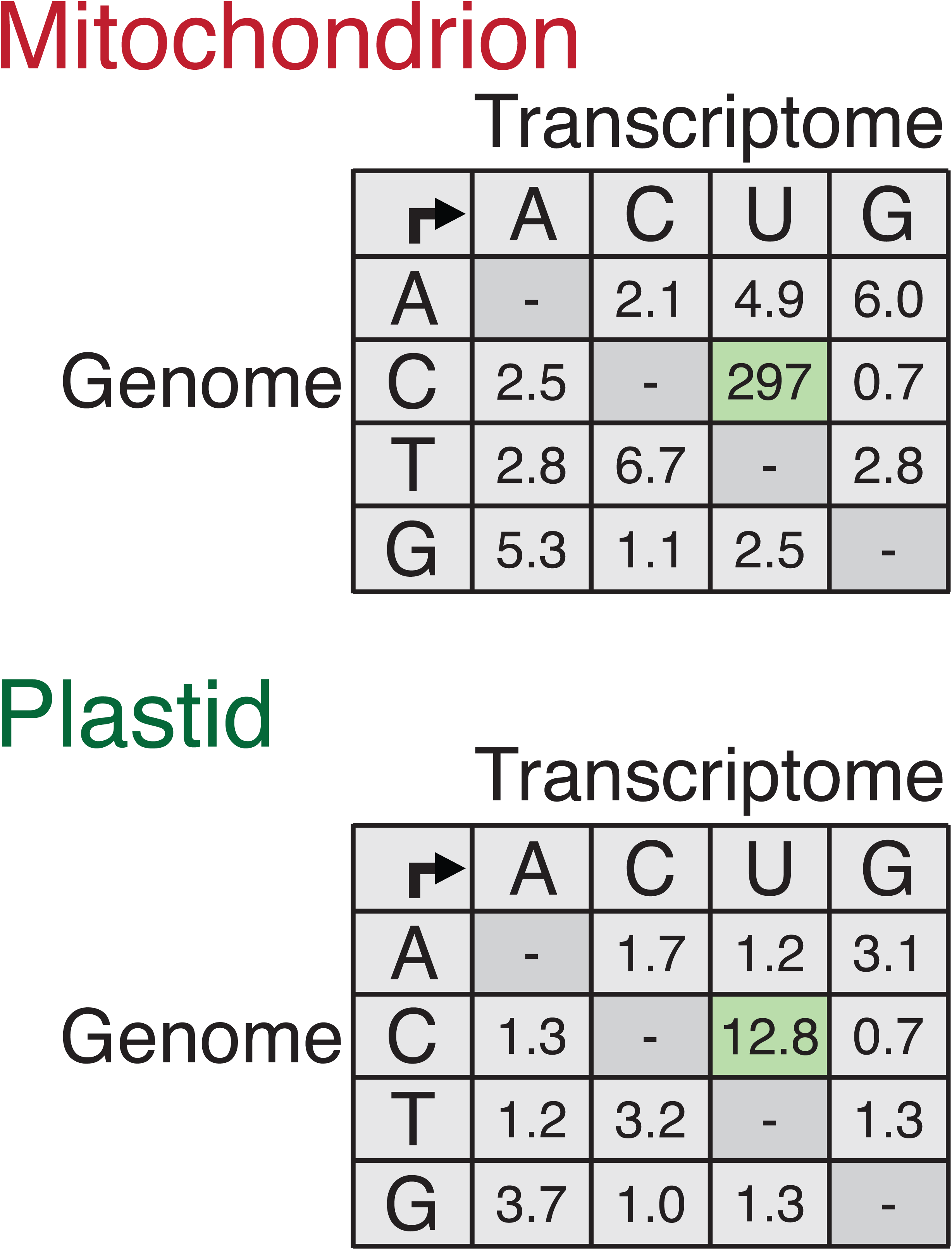
Comparison of differences between the genome and transcriptome of eight/fourteen species in the mitochondrial/plastid dataset, respectively. Values are number of sites (x10^3^) weighted by the number of parsiomony-informative sites across these species in each dataset and the average overlap between genome and transcriptome sequences (calculations documented in Table S8).

A complication of our simple approach is that the samples used to prepare genomes and transcriptomes for each species were not from the same genetic individual (Table S1, see also Bell et al., 2020) and so some of the differences likely represent within-species polymorphisms rather than RNA edits. It is reasonable to assume that all or nearly all the C-to-U differences constitute RNA edits rather than within-species genetic variation, as there are so many of them (see also Bell et al., 2020; Table S2, and Fig. 2 here). We also tentatively recognized U-to-C differences as possible edits here, as this type of edit does exist in other land plants (Knoop, 2023). The existence of such edits is unclear in angiosperms: Knoop (2023) noted that there has been sporadic reporting of U-to-C edits in angiosperms, which they were unable to replicate. The number of U-to-C changes is similar to other classes of genome-to-transcriptome differences noted here, see Table 2 (e.g., A-to-C differences). The latter cases almost certainly do not represent edits, as they would represent unknown forms of RNA editing in other land plants or other organisms (Knoop, 2023; though see Lonergan and Gray, 1993; Price and Gray, 1999 for examples in Amoebozoa tRNAs). Nonetheless, the impact of incorrectly recognizing a second class of RNA edits (U-to-C) on phylogenomic inference is likely to be small, as the number of inferred differences is not large (Fig. 2: Table S2).

### Phylogenetic analyses

We performed separate phylogenetic inferences of (i) a concatenated set of plastid genes (with and without putative edit sites removed); (ii) a concatenated mitochondrial gene set (with and without putative edit sites removed), and; (iii) a data set combining mitochondrial data (with putative edit sites removed) and plastid data (in this case without putative edit sites removed; as these edits had negligible effect on the inferred plastid phylogeny, see below). We performed maximum likelihood (ML)-based phylogenetic inference on these data sets using IQ-TREE v1.6.12 with default settings (20 initial best tree searches; Nguyen et al., 2015), partitioning the ML analyses by gene and codon (see Ross et al., 2016), and using the built-in IQ-TREE model tester to find the best models and partitioning scheme for each data set (Kalyaanamoorthy et al., 2017; models listed in Table S7). We inferred the gene by codon partitioning scheme and model testing on a smaller 80-taxon set due to computational limitations, using representative members from all major clades in the data set; we applied the resulting partitions to the full taxon set (see Tables S1, S7). We assessed branch support using 500 standard bootstrap replicates (Felsenstein, 1985), and mapped the bootstrap results onto the best tree using SumTrees (Sukumaran & Holder, 2010; Sukumaran & Holder, 2015). For the analysis of the plastid data with RNA-edit sites removed, we used the partition/model-testing results from the full plastid dataset, as plastid data have relatively few putative edits (see Figure 4).

## RESULTS

### Putative RNA-edit sites in mitochondrial and plastid data

We documented a total of 84– 734 genome-to-transcriptome nucleotide differences per each reference species examined, for their mitochondrial genomes (Table S2). Most of these differences (82–100% by species) correspond to C-to-U or U-to-C differences, with only a small number of other differences observed, i.e., zero to six differences for other nucleotide-nucleotide change classes, most of which correspond to transitions (Table S2). As non-C-to-U (or U-to-C) differences are not known as RNA edit types in other land plants (Knoop, 2023), they likely represent intraspecific variation (different genetic individuals were sampled for all genome and transcriptome pairs (Tables S1), although some may instead represent sequencing errors. For mitochondrial data, the lengths of genome-to-transcriptome overlaps within each reference species are summarized in Table S3 (by gene), along with counts of the total number of putative C-to-U RNA edits by gene (based on the overlapping regions in each reference species), counts of which C-to-U RNA edits are shared with those seen in at least one other reference taxon, and counts of “singleton” sites that are unique to each reference species. The within-gene alignment positions of each C-to-U RNA edit are also noted for each gene. Table S4 lists the C-to-U RNA edits that overlap for each pairwise reference-species comparison; counts and alignment positions are noted for each gene. For plastid data, we document the lengths of genome-to-transcriptome overlaps within each reference species, and counts of the total number of putative C-to-U RNA edits by gene (Table S9, paralleling Table S3 for mitochondrial data).

More broadly, across the reference species and regions where genome-to transcriptome comparisons were possible, we found a total of 849 and 19 putative C-to-U and U-to-C differences, respectively, for the mitochondrial dataset, and 289 and 72 putative C-to-U and U- to-C differences, respectively, for the plastid data set. Comparing these numbers between mitochondrial and plastid genomes is not straightforward, as the lengths of regions examined for genome-to-transcriptomes comparisons differ between the two genomes, and the two genomes evolve at substantially different rates (e.g., Wolfe et al., 1987). We therefore standardized these counts by adjusting for the length of overlapping genome-to-transcriptome regions (averaged across genes and taxa), and the number of parsimony-informative characters in alignments of the reference taxa. The resulting values are summarized in Fig. 2 for all classes of nucleotide-to-nucleotide differences, not only for the putative C-to-U RNA edits. It is evident that the C-to-U class of differences, expressed in relation to the total parsimony-informative variation per genome, is substantially the largest change class for plastid genomes—about an order of magnitude more frequent that other classes. The corresponding value of C-to-U differences is around two orders of magnitude higher for the mitochondrial genome compared to other change classes in that genome (Fig. 2, see also the summary in Table S2). This provides strong comparative evidence for the existence of C-to-U RNA edits in each organellar genome, and also documents the substantially higher level of editing in mitochondrial genomes compared to plastid genomes (at least an order of magnitude more C-to-U RNA edits per informative character are present for the mitochondrial data, based on these reference species).

### Accounting for RNA edit sites in phylogenomic inference

The plastid-based inferences appear to be essentially unaffected by inclusion vs. exclusion of RNA edit sites (compare tree figures and support values in Figs. S1–S4). We therefore only discuss the former analysis from here on for plastid data. By contrast, when putative RNA-edit sites are not accounted for in the mitochondrial phylogenomic analysis, all of the sampled orders, with the exception of Chloranthales and the eudicot orders, are recovered as non-monophyletic (Fig. 3, right-hand side, and see Figs. S5, S6). We also observed two or more sets of distantly related “sub-clades” for each of several orders (i.e., Austrobaileyales, Laurales, and Magnoliales), and the monocots— with sub-clade membership in each case seemingly determined by whether the component sequences were derived using transcriptomic or genomic data (note that the asterisks in Fig. 3 indicate taxa derived from transcriptome data). Several other major APG-based taxa (sensu APG, 2016) are also sub-divided into distantly related lineages in the resulting tree, including Piperales, monocots, and eudicots (although in these cases the resulting sub-clades did not exclusively represent genome vs. transcriptome-derived sequences; Fig. 3, right-hand tree). The order Nymphaeales is also divided into two distantly related sub-clades, with one sub-clade (comprising genome-derived sequences and a single transcriptome derived sequence) inferred to be sister to the rest of the angiosperms, and a second sub-cade inferred to be nested within a clade that includes monocots, Austrobaileyales, and *Amborella* (the latter species includes one sequence each derived from genome vs. transcriptome data). This sub-clade (comprising members of Nymphaeales, monocots, Austrobaileyales and *Amborella*) is in turn recovered as nested in a larger clade mainly comprising genome-derived sequences, which is in turn nested in a grade of Canellales taxa that include both transcriptome- vs. genome-derived sequences.

**Figure 3.**
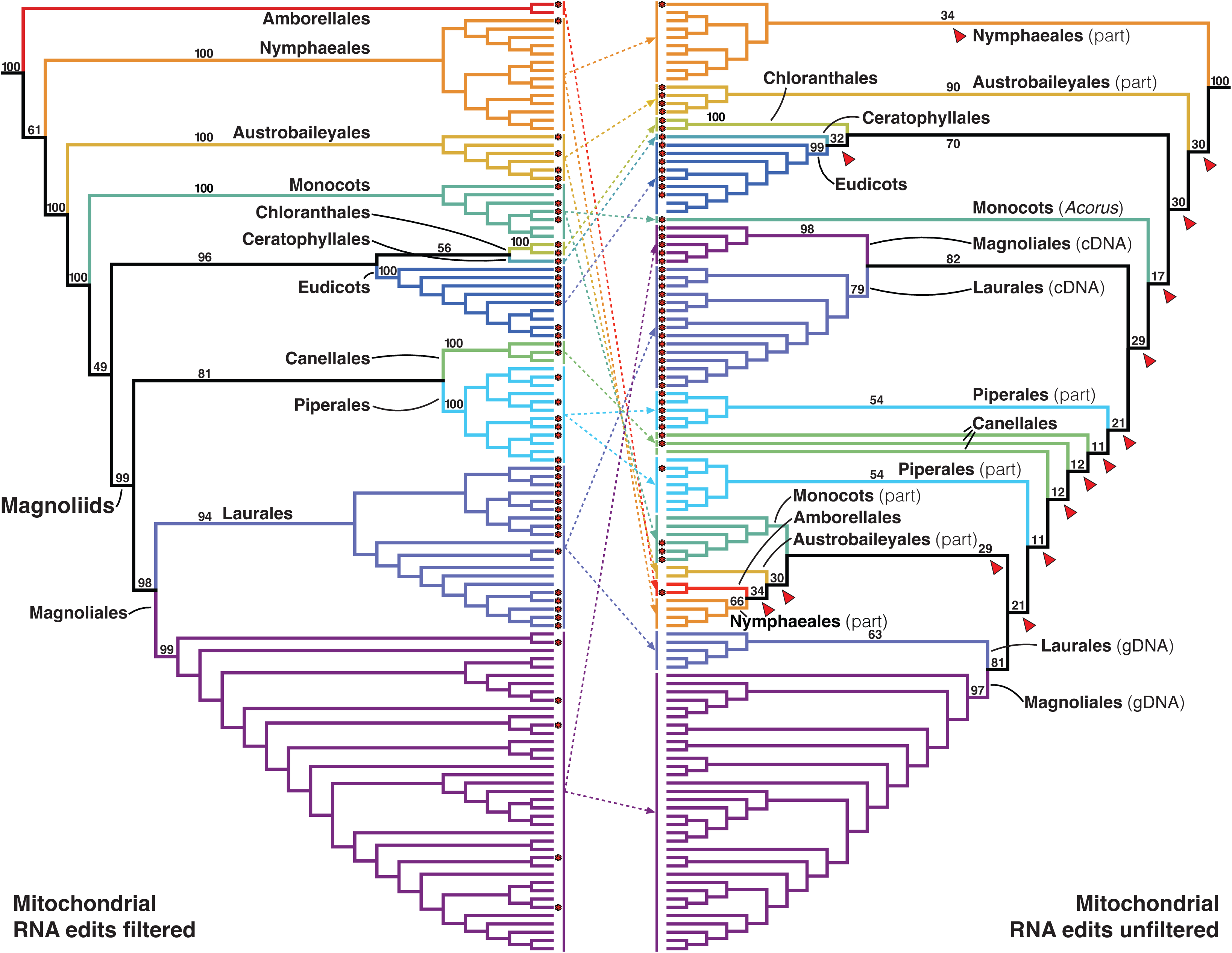
Disagreements in inferences of higher-order angiosperm relationships based on partitioned maximum likelihood (ML) analyses of mitochondrial genome data, in which putative RNA edit sites detected using a subset of reference taxa were filtered (left-hand tree, corresponding to the right-hand tree in Fig. 4), or included in analysis (right-hand tree here). Color-coded subclades and dotted lines indicate orders noted here (as currently recognized by APG 2016). Several clades composed of sequences solely derived from transcriptomes or genomes are indicated (cDNA; gDNA respectively). Species names, excluded for clarity here, are noted in Figs. S4–S8. Bootstrap support values from partitioned ML analyses are noted on a subset of branches (see Figs. S5–S8 for support values for other lineages).

These “broken” APG taxa, inferred when the RNA edit sites are included in analysis (Fig. 3, right-hand tree), conflict with well-accepted clades in plastid-based inferences (e.g., Fig. 4, left-hand tree), and are generally very poorly supported here by bootstrap analysis of the unfiltered mitochondrial data (Fig. 3, right-hand tree, and see Figs. S5, S6). In addition, branch support from the unfiltered mitochondrial data for well-accepted major clades observed in the plastid-based data set (e.g., Fig. 4, left-hand tree) is in general very poor (summarized in Table 1).

**Figure 4.**
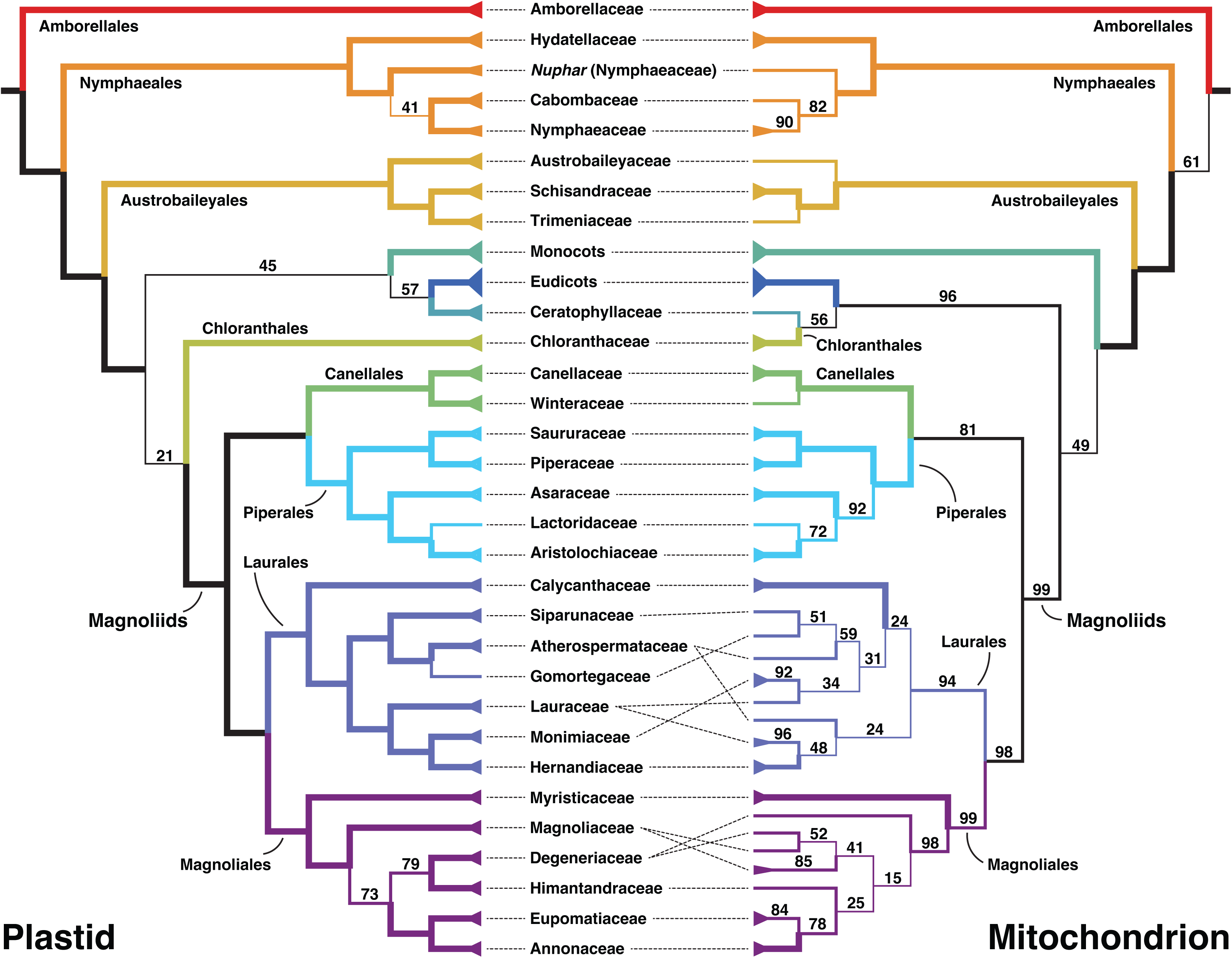
Inference of overall angiosperm relations based on plastid genome data (left side) and mitochondrial genome data corrected for putative RNA-edited sites (right side), both based on partitioned maximum likelihood (ML) analyses. Relationships shown focus on ANA-grade taxa (i.e., Amborellales, Nymphaeales and Austrobaileyales), and 20 magnoliid families in four orders, Ceratophyllales and Chloranthales, with distinct colors for individual orders. Individual families and the monocots, eudicots and are reduced here to single branches. Bootstrap support values are noted beside branches (thick branches indicate 100% support, regular branches indicate 70–99% support, and thin branches indicate <70% support). Dotted lines indicate some discrepancies in inferred higher-order relationships between plastid- and RNA-edit filtered mitochondrial-based inferences. Full details of relationships within each of these are presented in Figs. S1–S10.

**TABLE 1.**
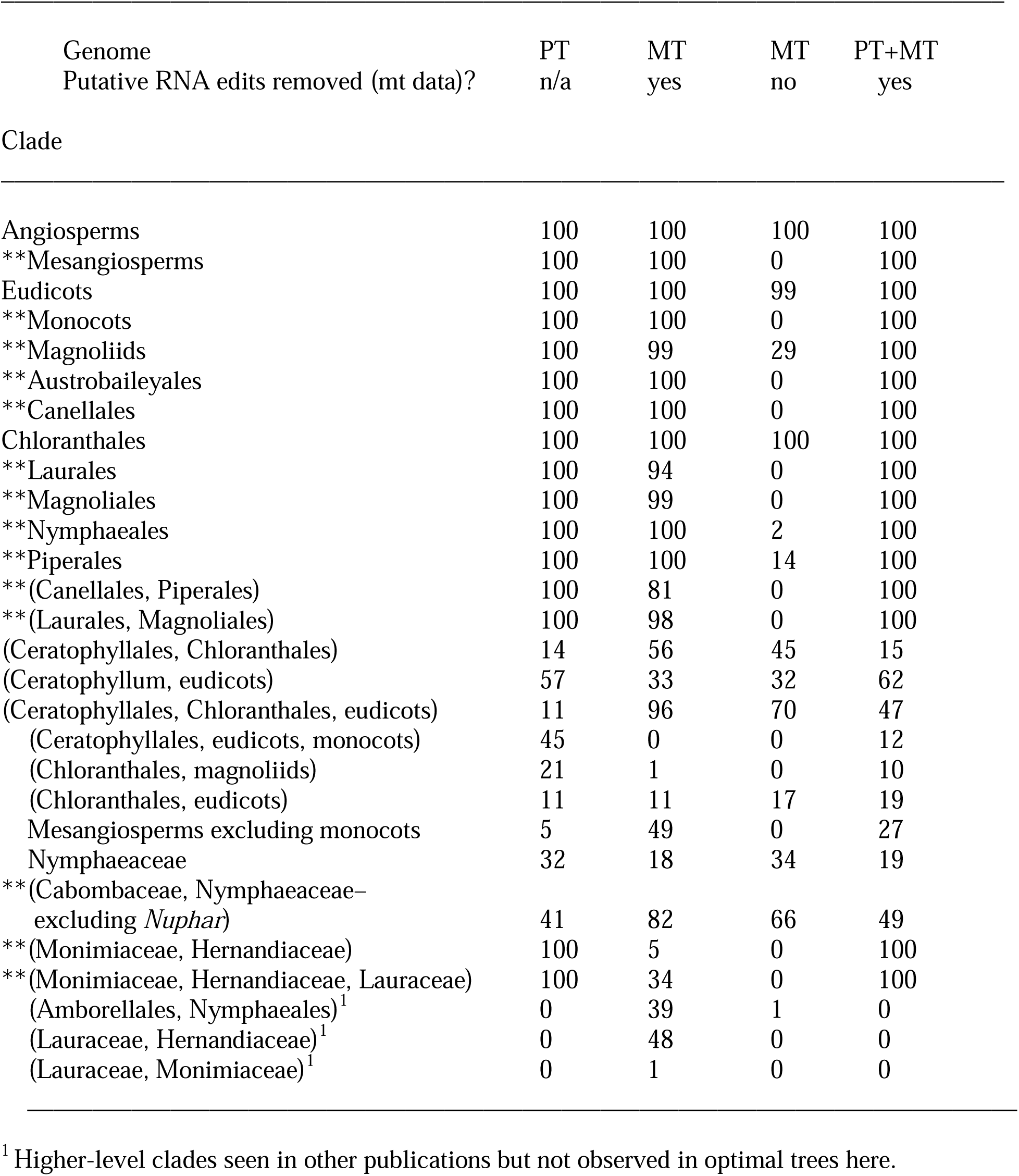
Likelihood bootstrap support values (percentages) for a selection of major angiosperm clades from plastid data, mitochondrial data and combined organellar data, illustrating strong divergence in a subset of mitochondrial results (clades marked with double asterisks, **) when putative RNA edits are included. Organellar genome (plastid = PT; mitochondrial = MT); see Table S5 for support values of additional major clades of interest.

Strikingly, however, the phylogenetic inferences based on filtered mitochondrial data (putative RNA edits sites removed) are largely congruent with APG-clades and our plastid-based inferences; they are also better supported (see Fig. 3, left-hand tree and right-hand tree; and Figs. S5–8 for full details). Thus, deleting the putative RNA edits from mitochondrial data appears to be sufficient to recover relationships that align with the plastid-based inferences. Most of the resulting higher-order clades are also well supported (Fig. 3) by the filtered mitochondrial data, with only weakly supported exceptions in Laurales and Magnoliales (Figs, 2, 3). Specifically, multiple families are recovered as non-monophyletic, but the resulting conflicts with monophyly of APG (2016) families are individually poorly supported; Fig. 3, right-hand tree, and see Figs. S7 and S8 for full details).

### Overview of higher-order relationships in angiosperms

From here on, we focus on results based on unfiltered plastid data and filtered mitochondrial data. The bulk of inferred higher-order relationships in angiosperms are both well-supported and congruent among all data sets, with major exceptions for major parts of the mesangiosperm “backbone” that also tend to be ambiguous in other published studies (Fig. 1), which we discuss below. Nearly all angiosperm families and orders according to APG (2016) (for those that are non-monotypic), and most larger clades recognized by APG, are well supported in analyses of plastid data and the combined organellar data (Fig. 4, left-hand tree; Figs. 5, S1, S2, S7–10). Exceptions include the family Nymphaeaceae, as a member of this family, *Nuphar*, is either weakly supported (in analyses of plastid data and combined organellar data), or moderately strongly supported (mitochondrial data) as sister to Cabombaceae and Nymphaeaceae (Figs. 4, 5, S2, S8, S10).

**Figure 5.**
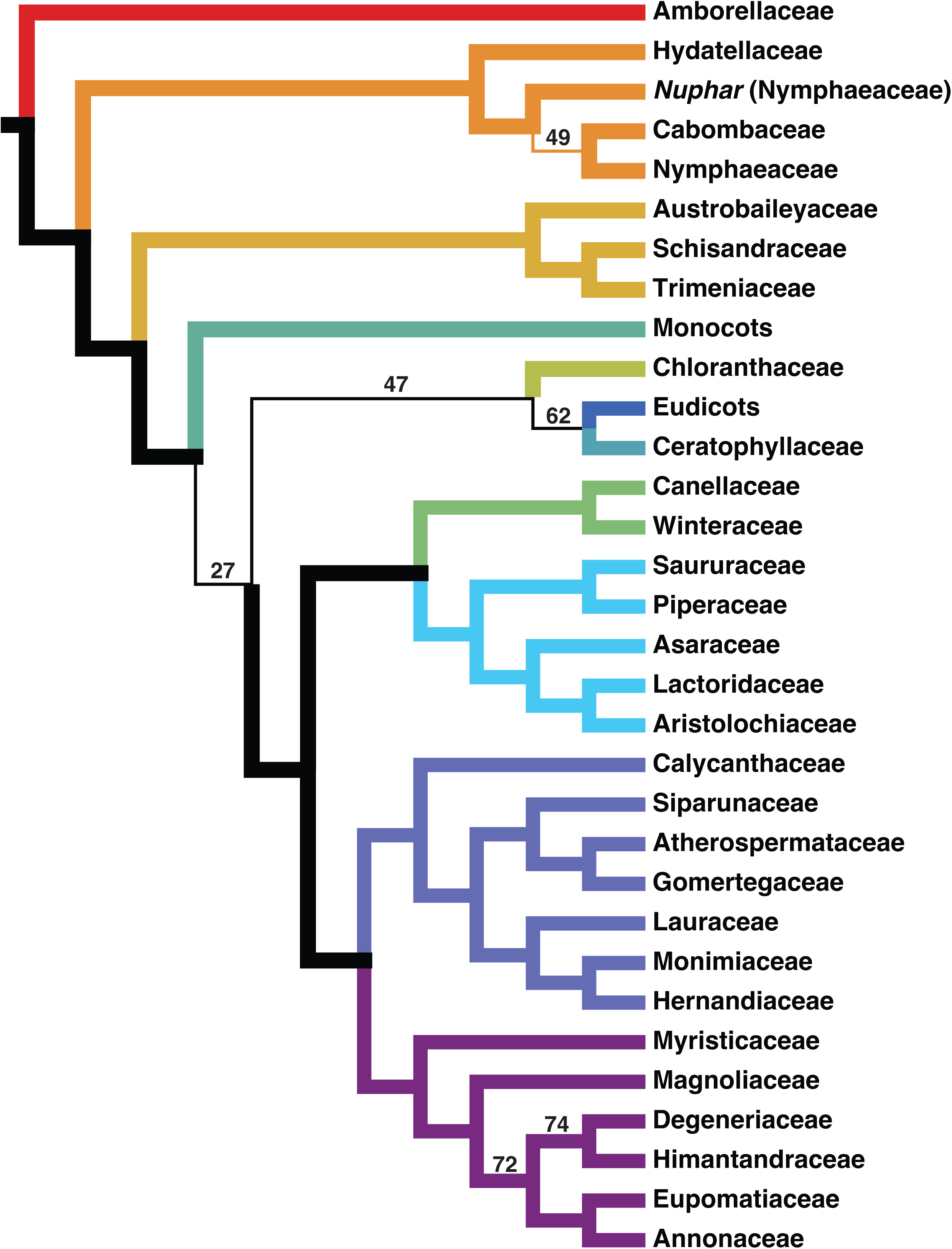
Inference of overall angiosperm relations based on a combined dataset of both plastid and RNA-edit filtered mitochondrial gene sets, based on partitioned maximum likelihood (ML) analysis. Coloring and branch thickness labels follow Figs. 3 and 4; the full tree is shown in Figs. S9 and S10.

Most relationships inferred among the ANA-grade and magnoliid orders, respectively, are well supported in analyses of the plastid data alone, the filtered mitochondrial data, and combined organellar data (Figs. 4, 5). However, analysis of the filtered mitochondrial data recovered only weak support for a magnoliid clade comprising Canellales and Piperales (81% bootstrap support), and only weakly supported a clade comprising all angiosperms except *Amborella* (only 61% bootstrap support). All three analyses strongly supported the monophyly of angiosperms.

Within the magnoliid orders, a major focus of our study, almost all inter-familial relationships are very well supported in the combined organellar analysis and by the plastid-only analysis (see thick branches in Fig. 4, left-hand tree and Fig. 5). The sole exceptions concern a group of four families in Magnoliales (i.e., Annonaceae, Degeneriaceae, Eupomatiaceae and Himantandraceae). A clade comprising these four families is moderately well supported in both analyses (72–73% support in the two analyses), as is a clade comprising Degeneriaceae and Himantandraceae (74–79% support). The relatively moderate support for these relationships may be because *Degeneria* (and thus the family as a whole) is represented here by one to few-gene data sets for two species, which have a very low number of plastid or mitochondrial genes in the alignments (two to four plastid genes, and one to two mitochondrial genes; Table S1). Degeneriaceae are also non-monophyletic in the filtered mitochondrial analysis (Fig. 4, right-hand tree; Fig. S7, S8), but this may be because the two *Degeneria* species are represented by non-overlapping genes for this data set (Table S6).

Relationships among the five major lineages in the mesangiosperms (Fig. 1) are largely uncertain here. For the plastid analysis, we recover a clade comprising eudicots and Ceratophyllales, with weak support (57% bootstrap support), a weakly supported clade comprising these taxa and monocots (45% bootstrap support), and a very poorly supported clade comprising Chloranthales and magnoliids (21% support; Fig. 4, left-hand side). For the filtered mitochondrial analysis, we recover a weakly supported clade comprising all mesangiosperms except monocots (49% bootstrap support), and a weakly supported clade comprising Ceratophyllales and Chloranthales (56% bootstrap support). However, the magnoliids are weakly inferred to be the sister-group to a strongly supported clade comprising eudicots, Ceratophyllales and Chloranthales (100% bootstrap support) by the filtered mitochondrial analysis. The combined organellar analysis recovers the same basic arrangement as the filtered mitochondrial data—i.e., (monocots, (magnoliids, (eudicots, (Ceratophyllales, Chloranthales))))—but with weak bootstrap support across all branches connecting these taxa (Fig. 5). Regardless of these differences, each component clade represented here by more than one species (i.e., Chloranthales, eudicots, monocots, and magnoliids) is also individually very well supported by all three data sets (Figs. 4, 5, S2, S8, S10).

We did not sample every eudicot and monocot order here, as this was not a focus of our study (16/49 and 7/12 orders sampled, respectively). However, we recovered strong support for most relationships at the order level and above, which we summarize here for the combined organellar tree (Figs. 5, S9, S10; the plastid tree has the same general topology with similar support values). Within monocots, we recover Acorales as the sister-group of all other monocots, followed by Alismatales, Dioscoreales, Liliales, Asparagales, and Arecales-Poales as successive sets of sister-groups, with 100% support for most of these arrangements (an exception is a branch supporting the position of Dioscoreales, which has 67% support). Within the eudicots, we recover Ranunculales as the sister group of the remaining eudicots, followed by Proteales, Trochodendrales and Buxales as successive sister-groups of the core eudicots, all with strong support (98–100%). Within the core eudicots, we recover Gunnerales as the sister group of the remaining taxa, with 100% support. Within the superasterids, we found Caryophyllales to be the sister group of the asterids, with Asterales, Gentianales, and Lamiales-Solanales forming succesively sister-group relationships within the asterids, all with with 100% support (apart for a sister-group relationship between Lamiales and Solanales, which has 60% support). Within the superrosids, we recover Saxifragales and Vitales as sister groups with 66% support; this clade is in turn sister to the rosids, with 99% support. Within rosids, we recover Fabales (with 100% support) as the sister group of a clade comprising Geraniales, Brassicales, and Malvales which has 80% support; the latter two orders are also inferred to be sister groups, with 100% bootstrap support.

We included a relatively large sampling of gymnosperms as outgroups. We arbitrarily rooted the seed plants such that gymnosperms are sister to angiosperms (e.g., Fig. S1**–**10), as we did not include more distant outgroups (a caveat of our reporting here is therefore that the monophyly statements in this section depend on where the true seed-plant root belongs). For all analyses, both angiosperms and gymnosperms are strongly supported as distinct clades (100% support; Figs. 4, 5). For both the plastid-only and combined organellar datasets, we found 81% and 54% support, respectively, for placing Gnetales as the sister group of cuppresophyte conifers (i.e., Araucariaceae, Cephalotaxaceae, Cupressaceae, Podocarpaceae, Sciadopityaceae and Taxaceae; which also together form a clade with 100% support in both analyses), the so-called “GneCup” relationship (Figs. S2, S10). By contrast, we found 89% support for a relationship with Gnetales as the sister group of Pinaceae (with 100% support; the “GnePine” relationship) for the filtered mitochondrial data set (Fig. S8). For the plastid dataset, we recovered Cycadales as the sister group of the rest of the gymnosperms (with 100% support), with *Ginkgo* then the sister group of the GneCup clade, with 87% support (Fig. S2). By contrast, in the analyses of the filtered mitochondrial and combined data sets, we recovered a clade comprising only Cycadales and *Ginkgo* (with 85% and 70% support, respectively); this Cycadales-Ginkgoales clade was then recovered as the sister group of a clade comprising conifers and Gnetales (with 100% support in both analyses; Figs. S8, S10).

### Relationships within two densely sampled families

Most of the samples in Hydatellaceae are represented by whole plastid genomes, and by at least 50% of the full set of mitochondrial genes (Table S6). A few taxa are instead represented by small gene sets (four of seventeen terminals for plastid data; a separate three of nine terminals for mitochondrial data). Our plastid and combined organellar analyses recover the same topology (Figs. S2, S10), with mostly similar support at all branches. For the filtered mitochondrial analysis (Fig. S8) we infer the same underlying set of relationships for terminals in common. We therefore refer to the combined analysis here for simplicity (see Fig. S2 for branch support in the plastid analysis). Relationships between all four sections of *Trithuria* have 100% support (see Fig. S10), and support for relationships within each section are almost very well supported (all but one branch has 91– 100% support; Fig. S10). The topology found here for the section *Trithuria* is similar to the one found using four plastid markers by Iles et al. (2012), but resolves several polytomies in the latter. Within section *Trithuria* we find strong support for non-monophyly of *T. bibracteata*, as *T. submersa* is well supported as being nested within the former species (with 91–92% support for relevant branches; Fig. S10). Within section *Hydatella*, we recover *T. austinensis* as the sister-group of the rest of the section with strong support (100%), generally with strong support for the remaining relationships. The clade that includes *T. australis*, *T. fitzgeraldii* and the specimen labeled *Trithuria* aff. *australis* has one or more cryptic species that still require proper naming (Sokoloff et al., 2019).

Within Trimeniaceae, only two species (*T. moorei* and *T. nukuhivensis*) are represented by plastid genomes (Table S6), and the mitochondrial genome was sequenced for only one taxon, *T. nukuhivensis*. The plastid data set is otherwise represented by few-gene data derived from Sanger-based sequencing (i.e., eight genes in total; Table S6). Here we recover two main subclades in both the plastid and combined organellar analyses, one containing two lianas (*Trimenia moorei* and *T. macrura*), which is then the sister group of *T. neocaledonica* (all of these relationships have 100% support in both analyses; Figs. S2, S10). The other major subclade comprises (in terms of successive sister-group relationships): *T. papuana,* then *T. nukuhivensis*, and then a small clade comprising *T. marquesensis* and *T. weinmannifolia* (with at least 95% bootstrap support for all relevant branchs in both analyses; Figs. S2, S10).

## DISCUSSION

A major aim of our study was to explore the effect of RNA editing on analyses, particularly those based on mitochondrial data, where the impact appears to be extreme (Fig. 3). We identified RNA editing in the mitochondrial genome as having a major misleading effect on phylogenetic inferences when using mixed sources of genomic and transcriptomic data, as predicted by Bowe and DePamphilis (1996; e.g., Fig. 1). Fortunately, these putative edit sites, which are typically shared across multiple taxa (Table S3, S4), are readily uncovered and accounted for, using reference taxon comparisons, addressing the second major goal of our study. The resulting phylogenetic inferences are largely congruent with other published plastid and nuclear phylogenies (e.g., One Thousand Plant Transcriptomes Initiative, 2019; Li et al., 2021; Zuntini et al., 2024). Our results should therefore mitigate fears that the mitochondrial genome is not a good source of phylogenetic information for higher-order plant phylogenetic inferences, in line with other recent phylogenomic studies of this organelle (e.g., Bell et al., 2020; Dong et al., 2022; Lin et al., 2022; Soto Gomez et al., 2020; Xue et al., 2022; Klimpert et al., 2022).

A third goal of our study was to perform combined organellar analysis (after accounting for RNA edits sites in the mitochondrial data). The resulting analysis (Fig. 5) mostly matches the plastid-based phylogenetic tree, and the filtered mitochondrial tree. It provides a comprehensive view of organellar genome phylogeny, providing a useful point of comparison for future more densely sampled nuclear-based studies. In several under-sampled lineages (e.g., Degeneriaceae, Hydatellaceae, Trimeniaceae) we also explored the effect of including few-gene datasets in combination with whole organellar gene sets, which appears to work well for Hydatellaceae, and Trimeniaceae, but less well for Degeneriaceae (in this family there there were very few available genes, and they did not fully overlap between taxa for mitochondrial data).

### Mitochondrial RNA editing

Mitochondrial genomes provide a potentially rich (and in many ways largely untapped) resource for understanding the early angiosperm radiation (see Xue et al., 2022 for a recent study that used them effectively for this question). However, phylogenomic studies of higher-order plant relationships have generally not included data from this genome (e.g., the One Thousand Plant Transcriptomes Initiative, 2019; Li et al., 2019, 2021; Zuntini et al., 2024). This may in part be due to concerns about the very slow rate of evolution of individual mitochondrial genes in most plant lineages, which has been recognized since the early days of molecular systematics (e.g., Wolfe et al., 1987). Nonetheless, the very slow rate of mitochondrial genome evolution may not be a problem *per se*, as multiple recent phylogenomic studies have shown that by combining the full array of protein-coding sequences from this genome, we can obtain well supported phylogenomic inferences. In effect, the combined mitochondrial gene set collectively provides sufficient variation for robust inference of high-order phylogenetic relationships among lineages (e.g., Bell et al., 2020; Dong et al., 2022; Lin et al., 2022; Soto Gomez et al., 2020; Xue et al., 2022; Klimpert et al., 2022).

The relatively limited use of mitochondrial phylogenomic data to date also likely reflects a well-founded concern that RNA editing can have a misleading effect on phylogenetic inference when genomic and transcriptomic data are combined (e.g., Bowe and DePamphilis, 1996). Xue et al. (2022) examined the early angiosperm radiation using mitochondrial genome data, but their study did not mix genomic and transcriptomic data (note that they also used a substantially smaller sampling of the major lineages included here). We know that RNA editing in mitochondrial genes can be a substantial problem in large-scale phylogenetic studies that include data derived from both transcriptomes and genomes, a phenomenon addressed only rarely in land plants (e.g, Bowe and DePamphilis, 1996; Qiu et al. 2006, 2010; Bell et al., 2020; Dong et al., 2022). Here we compared mitochondrial analyses in which putative RNA edit sites were either removed or not, which we identified using a core set of reference taxa with parallel genome and transcriptome data. Our results demonstrate that accounting for RNA edit sites in this way is highly effective in correctly anomalous relationships or very poor bootstrap support in mixed analysis of genomic and transcriptomic data (e.g., Fig. 3, 4).

This approach is relatively straightforward to do using reference taxa spanning the lineages of interest. When we did not correct for putative RNA edit sites, the resulting phylogenomic inference displays numerous highly anomalous—and poorly supported— relationships compared to our current understanding of angiosperm phylogeny (e.g., One Thousand Plant Transcriptomes Initiative, 2019; Xue et al., 2022; Li et al., 2021; Zuntini et al., 2024) (Fig. 3, right-hand tree). This is despite only ∼1.8% of the total alignment length having predicted RNA edit sites (Fig. 2, Table S6, also see Supplementary Material). Here we also removed putative U-to-C RNA edits, an RNA-edit class that exists in several land-plant lineages (hornworts, lycophytes and ferns; Knoop, 2023), although not unambiguously reported in any angiosperm organellar genomes to date (Knoop, 2023, and see Ruchika et al., 2021, for evidence of U-to-C edits in nuclear genes). We included clades beyond those that have been best-studied for RNA editing (i.e., eudicots and monocots) (Knoop 2023). It is not inconceivable that this RNA-edit class exists but is unreported in angiosperms. Regardless, filtering for this possible RNA-edit class likely had very little effect on the resulting phylogenetic inferences, as only 19 putative U-to-C edits were found in the mitochondrial dataset (representing 0.04% of the total alignment length).

Strikingly, removing all putative RNA edits sites from the mitochondrial data sets greatly reduces or removes these numerous highly anomalous relationships (Fig. 3; left-hand vs. right-hand trees), resulting in a mitochondrial phylogenomic inference that closely matches the one based on plastid data (summarized in Fig. 4 here). This updated mitochondrial-based inference also has substantially improved branch support (Fig. 3). Bell et al. (2020) found similar levels of RNA editing in their bryophyte-focused study (∼2% of total alignment length), although their results showed strongly supported local conflicts between inferences based on alignments with RNA edits removed vs. retained dataset, whereas our study depicts very low support for most higher-order relationships when putative RNA edit sites are not filtered (Fig. 3, right-hand tree).

Our mitochondrial RNA edit predictions are based on a core set of genome/transcriptome comparisons (only eight species from across a larger sampling of 141 taxa; Table S1), but the efficacy of our approach (e.g., Fig. 3) supports the idea that many of these putative edits are shared across angiosperm phylogeny as a whole (see also Table S3). The impact of RNA-edit sites on phylogenetic or phylogenomic inference is consistent with early phylogenetic findings on land plants by Bowe and DePamphilis (1996) based on phylogenetic evidence from only two genes, and with two more recent studies of overall land-plant and liverwort phylogeny using organellar phylogenomic data sets (Bell et al., 2020; Dong et al., 2022). Indeed, recent studies have demonstrated the utility of mitochondrial genome data for resolving difficult-to-place heterotrophic lineages in angiosperm phylogeny (Lin et al., 2022), and in reconstructing overall land-plant phylogeny (Bell et al., 2020). Mitochondrial RNA-edit sites did not appear to substantially impact phylogenetic inference in the study by Lin et al. (2022), but this is not particularly surprising as their alignments did not include mixed genomic and transcriptomic sequences.

Why is the impact of RNA editing on phylogenetic inference substantially larger for mitochondrial genomes than plastid genomes (Figs. 2, S5**–**8)? First, there is a nearly order of magnitude more RNA edits in their protein-coding genes per nucleotide (1.8% vs 0.4% of all sites in the mitochondrial vs plastid dataset, respectively; e.g., Takenaka et al., 2013; Fig. 2, Table S2, Table S3, S9). In addition, mitochondrial genes also evolve very slowly (Wolfe et al., 1987), and they have fewer protein-coding genes (typically ∼42 genes; with 10,927 parsimony-informative sites in our alignment) than plastid genomes (∼79 genes; with 35,171 parsimony-informative sites in our alignment). This may make phylogenomic analysis based on mitochondrial data more prone to any distorting effect of RNA editing, compared to inferences based on the more rapidly evolving and larger plastid-based analyses.

### Other key phylogenetic findings

Our study was focused on documenting and accounting for the effect of RNA editing on phylogenomic inference, but we also performed the first combined phylogenomic study of the two organellar genomes (Figs. 3, 4, 5, S1-10; at this taxonomic scale) and included multiple new taxa (Table S1). The positions of the newly added taxa within families and clades are highlighted in Fig. S1, S3, S5, S7, S9 (see red asterisks) and generally conform with other studies (e.g., Massoni et al., 2014, Helmstetter et al., 2025). We also resolved relationships among three families in Laurales here (i.e., Hernandiaceae, Lauraceae and Monimiacieae) that have long been controversial (e.g., Qiu et al., 1999, 2000, 2005, 2006, 2010; Renner, 1999, 2004; Renner and Chanderbali, 2000; Savolainen et al., 2000; Hilu et al., 2003; Zanis et al., 2003; Soltis et al., 2011; Massoni et al., 2014). Within Magnoliales, we were able to resolve Degeneriaceae as the sister group of Himantandraceae with moderate support in the plastid and combined analyses (Fig. 4, left-hand tree; Fig. 5); the relationship between these families has proved hard to resolve with good support (e.g., Sauquet et al., 2003; Massoni et al., 2014). The relationship recovered here aligns with Helmstetter et al. (2025). However, both species of *Degeneria* included here were represented in both organellar data sets by only a few genes, and so this taxon will likely least require full organellar genomes to resolve its position with more clarity.

We were unable to satisfactorily resolve a long-standing controversy over the relative relationships among the five major mesangiosperm lineages (Fig. 1), although the filtered mitochondrial data set recovered a strongly supported clade comprising Ceratophyllales-Chloranthales-eudicots (Fig. 4, right-hand-side, Fig. S8). However, the combined data set recovered the same clade but with reduced support (Fig. 5, S10), perhaps hinting at additional residual but cryptic conflict in phylogenetic signal for this relationship, between the two organellar genomes. Notably, the filtered mitochondrial data set also did not recover strong support for the position of *Amborella* as the sister group of other angiosperms (Fig. 4, right-hand-side, Fig. S8), and there is also strong conflict concerning the relationships among the major groups of gymnosperms (Fig. S7, S8), consistent with similar conflicts in many other studies (Chaw et al., 1997, 2000; Bowe et al., 2000; Schmidt and Schneider-Poetsch 2002; Graham and Iles, 2009; Ran et al., 2010, 2018; Wu et al., 2011; Lu et al., 2014; Stull et al., 2021; Liu et al., 2022). The filtered mitochondrial data recover, although with low support, a sister-group relationship between two species-poor and phylogenetic “wildcard” mesangiosperm lineages, Ceratophyllales and Chloranthales. This grouping was also found in a few earlier molecular studies involving plastid (Antonov et al., 2000), nuclear (Zeng et al., 2014) and mitochondrial data (Xue et al., 2020). It fits well with morphological data (Endress and Doyle, 2009, 2015; Doyle and Endress, 2018; Sokoloff et al., 2022). Cretaceous fossils potentially related to both Ceratophylles and Chloranthales are important in this context (Kvaček et al., 2016, Doyle and Endress, 2018). Even if the Ceratophyllales-Chloranthales clade is an artefact of analyses, these fossils, especially *Canrightia* (Friis and Pedersen, 2011), may still be instructive in inferring ancestral morphology of lager clades such as the Ceratophyllales-Chloranthales-eudicots clade found in the present study.

Several families in the ANA grade that have received little attention using phylogenomic data to date, were included here with mixed genomic and Sanger-based few-gene data sets. The resulting inferences are consistent with but more resolved than previously published studies for Hydatellaceae (Iles et al., 2012; Marques et al., 2016; Sokoloff et al., 2019), and they also strongly support incongruence found between plastid data in general and nuclear (ITS) data for this family (Iles et al., 2012). In particular, the eastern group of *T. submersa* was inferred to be the sister-group of the rest of section *Trithuria* for the nuclear data (Iles et al., 2012), which conflicts with the plastid findings (e.g., we recover a sister-group relationship between eastern and western populations of this species here, e.g., Fig. S2). Reticulate evolution and polyploidy apparently played role in the evolution of yet unnamed cryptic species in the section *Trithuria* (Kynast et al., 2014; Marques et al., 2016). We allow for the first time to inferred a well-supported set of relationships for Trimeniaceae, where we observed the lianas (sometimes recognized as a separate genus, *Piptocalyx* Oliv. ex Benth.) as nested within *Trimenia* (Fig. S1, S2, S9, S10). Thus, our study also demonstrates that combining legacy Sanger-based data sets with phylogenomic data is feasible, showing how genomic data can be readily connected to the wealth of older molecular systematic studies that have defined the field over the last three decades and more.

### Conclusion

Phylogenomic mitochondrial data sets that are derived from mixed sources (genomic and transcriptomic) can produce anomalous and poorly supported inferences of higher-order angiosperm relationships. However, this systematic bias can be readily handled by filtering for problematic RNA edit sites in mitochondrial data. We can predict these sites using reference taxa for which we have both genomes and transcriptomes. The resulting inferences generally align well with several decades of evidence on these relationships. We used only a small set of representative reference taxa to identify problematic RNA edit sites, and it will be undoubtedly be useful to include more reference taxa for inferring these sites in future, as genomic and transcriptomic data sets become increasingly ubiquitous, and additional organellar genomes continue to be produced incidentally as byproducts in other genomic studies (e.g., Bell et al., 2020; Baker et al., 2021). We included a large number of previously unsampled lineages that define the early angiosperm radiation for both organellar genomes, and our study includes by far the largest sampling for these lineages attempted to date using mitochondrial genome data alone. Our analyses generally revealed congruent inferences between organellar data sets, when mitochondrial genomes are filtered for RNA edits. Nonetheless, we were unable to resolve relationships among the five major mesangiosperm lineages using combined organellar data alone, although we found strong support for a clade comprising Ceratophyllales, Chloranthales and eudicots using the filtered mitochondrial data, which warrants further investigation.

## Acknowledgments

The authors would like to thank the following organizations and individuals: Nate Klimpert (University of British Columbia) for help with scripts and computational issues related to analyses, and Laetitia Carrive (Université Paris-Sud) for assistance with lab work associated with Annonaceae genome skims. We thank Department of Parks and Wildlife Western Australia; Parks and Wildlife Commission, Northern Territory and the Director of National Parks, Australian Government for permission to collect material of Hydatellaceae used here, and the Australian Geographic Society for funding field travel (for TM). Analyses were completed in part using servers from the Digital Research Alliance of Canada (alliancecan.ca); some analyses used GNU Parallel (Ole 2023). We thank the German Academic Exchange Service (DAAD) for funds to S.W. for exchange between Germany and Canada (PPP Canada), the academic exchange office of TU Dresden and the Leonardo office Dresden, and Erasmus+ KA107 action for travel funds and mobility organization This research was also supported by an Agence Nationale de la Recherche grant (number ANR-12-JVS7-0015-01) to HS, and by a Natural Sciences and Engineering Research Council of Canada (NSERC) Discovery grant to S.W.G.

## Author Contributions

I.M., B.T.S., W.I., M.M., S.L., T.L.P.C., and H.S. performed DNA sequencing. D.L., T.F., S.M., and S.W. provided material; W.K.G., M.J. and H.S. assembled genome skim data and extracted gene sets. W.K.G. performed analyses. W.K.G., and R.V. developed scripts; W.K.G., I.M., and S.W.G. conceived the study and wrote the paper, with contributions from all authors.

## APPENDICES

Fig. S1. Phylogram of optimal plastid-based phylogenomic tree of angiosperms and relatives using a partitioned DNA-based maximum likelihood (ML) analysis (gene x codon partitioning scheme, see text). The sampling has a focus on ANA-grade and magnoliid taxa (seed plants arbitrarily rooted between angiosperms and gymnosperm outgroups). Species new to this study are indicated with a red asterisk (with corresponding voucher information; Table S1). Previously published species indicated with GenBank numbers or a four-letter code from One Thousand Plant Transcriptome Initiative (2019); several published sequences lacking GenBank numbers are indicated with author names from the corresponding publication (Hoekstra et al. 2017, Li et al. 2019 & 2021, Rossetto et al. 2015), followed by associated voucher numbers. Species represented by only a few genes are indicated with a dagger (with voucher or GenBank accession noted for each gene). Otherwise, species represented by pooled data from different specimens (from previously published studies or the current study) are indicated with multiple voucher numbers (Table S1). Species with a "+" indicate multiple Genbank accessions from the same specimen (see Table S1 for a complete list). The scale bar indicates estimated number of substitutions per site.

Fig. S2. Cladogram of optimal plastid phylogenomic tree from Fig. S1, with associated bootstrap support values.

Fig. S3. Phylogram of optimal plastid phylogenomic tree with all C-to-U and U-to-C genomic vs. transcriptomic differences removed from consideration (DNA-based ML analysis with a gene x codon partitioning scheme, see text). Scale bar indicates estimated number of substitutions per site. See Fig. S1 for explanation of symbols and provenance information.

Fig. S4. Cladogram of optimal plastid phylogenomic tree in Fig. S3, with associated bootstrap support values.

Fig S5. Phylogram of the optimal mitochondrial phylogenomic tree with all C-to-U and U-to-C genomic vs. transcriptomic differences included (DNA-based ML analysis with a gene x codon partitioning scheme, see text). Scale bar indicates estimated number of substitutions per site. See Fig. S1 for explanation of symbols and provenance information.

Fig. S6. Cladogram of the optimal mitochondrial phylogenomic tree in Fig. S5, with associated bootstrap support values.

Fig S7. Phylogram of optimal mitochondrial phylogenomic tree with all C-to-U and U-to-C genomic vs. transcriptomic differences removed from consideration (DNA-based ML analysis with a gene x codon partitioning scheme, see text). Scale bar indicates estimated number of substitutions per site. See Fig. S1 for explanation of symbols and provenance information.

Fig. S8. Cladogram of the optimal mitochondrial phylogenomic tree in Fig. S7, with associated bootstrap support values.

Fig S9. Phylogram of the optimal phylogenomic tree inferred from combined organellar (plastid and mitochondrial) data, with all mitochondrial C-to-U and U-to-C genomic vs. transcriptome differences removed from consideration (DNA-based ML analysis with a gene x codon partitioning scheme, see text). Scale bar indicates estimated number of substitutions per site. Blue diamonds indicate species with plastid data only. See Fig. S1 for explanation of symbols and provenance information.

Fig. S10. Cladogram of the optimal combined organellar tree inferred in Fig. S9, with associated bootstrap support values.

Table S1. List of taxa sampled in this study.

Table S2 Nucleotide variation between species with both genomic and transcriptomic data.

Table S3. Lengths of recovered sequences for each species and gene, including the genome/transcriptome overlap within species, and counts and lists of predicted RNA edit sites per taxon, and the fraction of those shared in overlapping regions with other species (see Table S4).

Table S4. List of the positions in the full mitochondrial matrix that overlap between species.

Table S5. Bootstrap support for major clades in angiosperm phylogeny (excluding those within monocots and eudicots).

Table S6. List of genes sampled in plastid and mitochondrial datasets.

Table S7. Models inferred from IQ-TREE model-tester.

Table S8. Counts of site changes from genome to transcriptome in fourteen/eight species examined in the plastid/mitchondrial data, respectively.

Table S9. Lengths of recovered plastid sequences for each species and gene, including the genome/transcriptome overlap within species, counts and lists of predicted RNA edit sites per taxon, and the fraction of those shared in overlapping regions with other species.

