## Supplementary figures and images for "Organellar data sets confirm overall angiosperm relationships if problematic RNA-edit sites are accounted for in mitochondrial genomes"

### Fig. S1

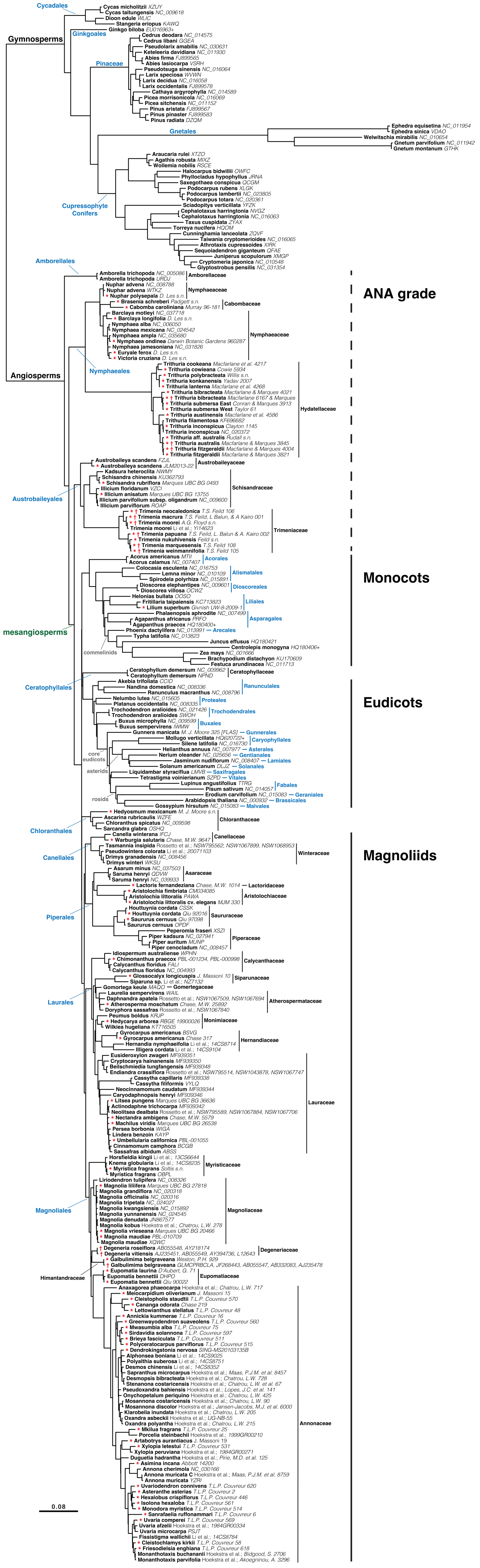

### Fig. S2

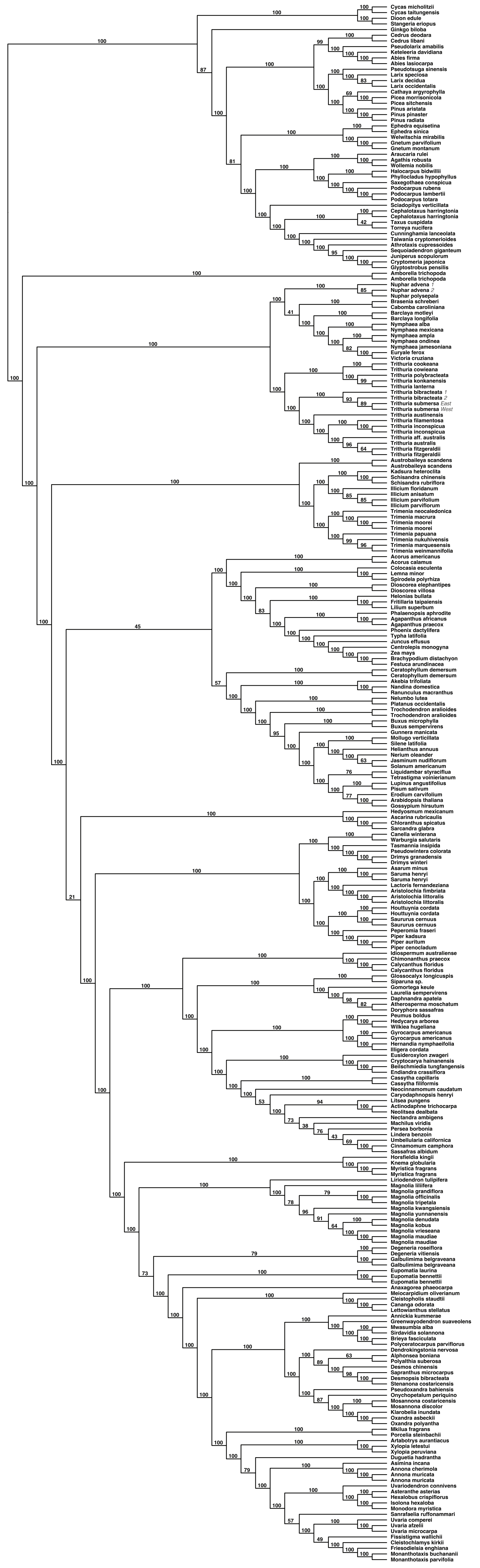

### Fig. S3

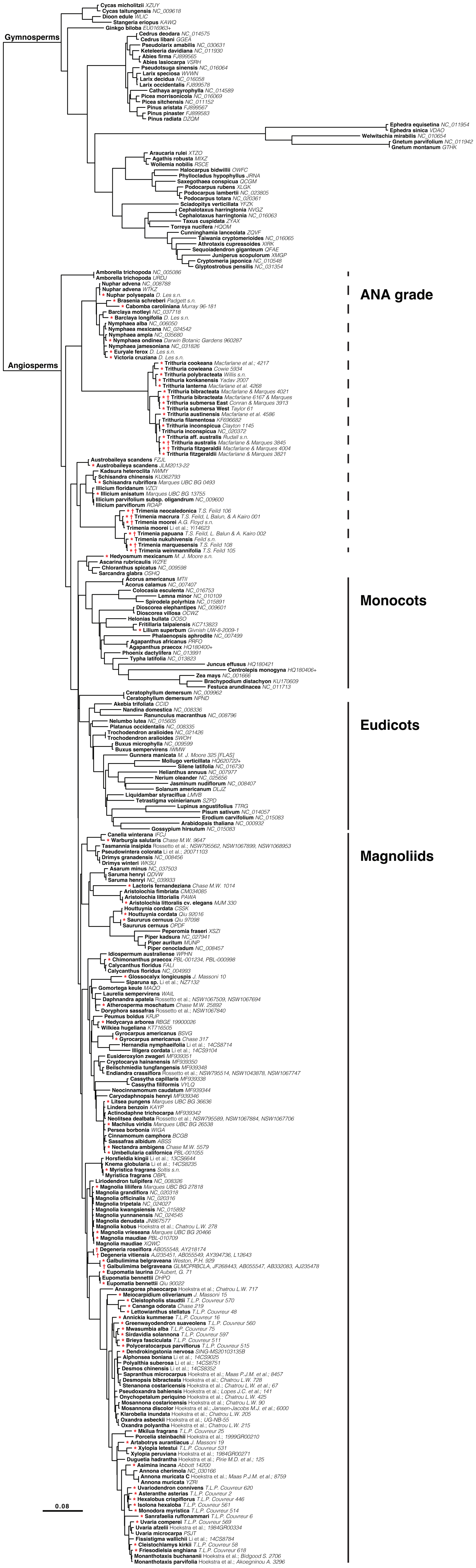

### Fig. S4

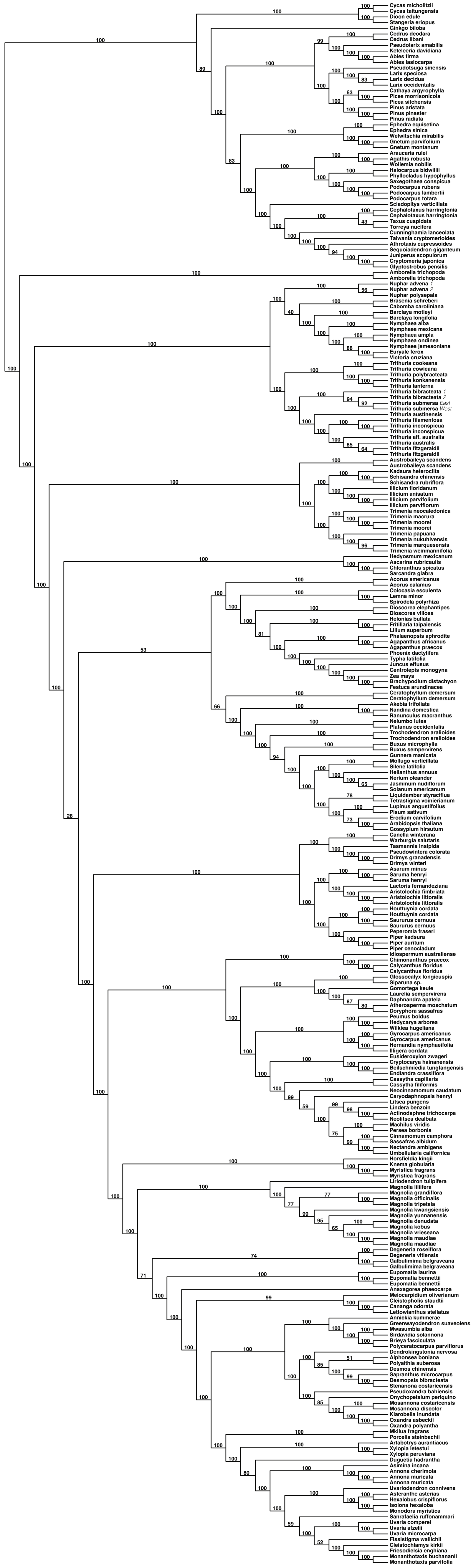

### Fig. S5

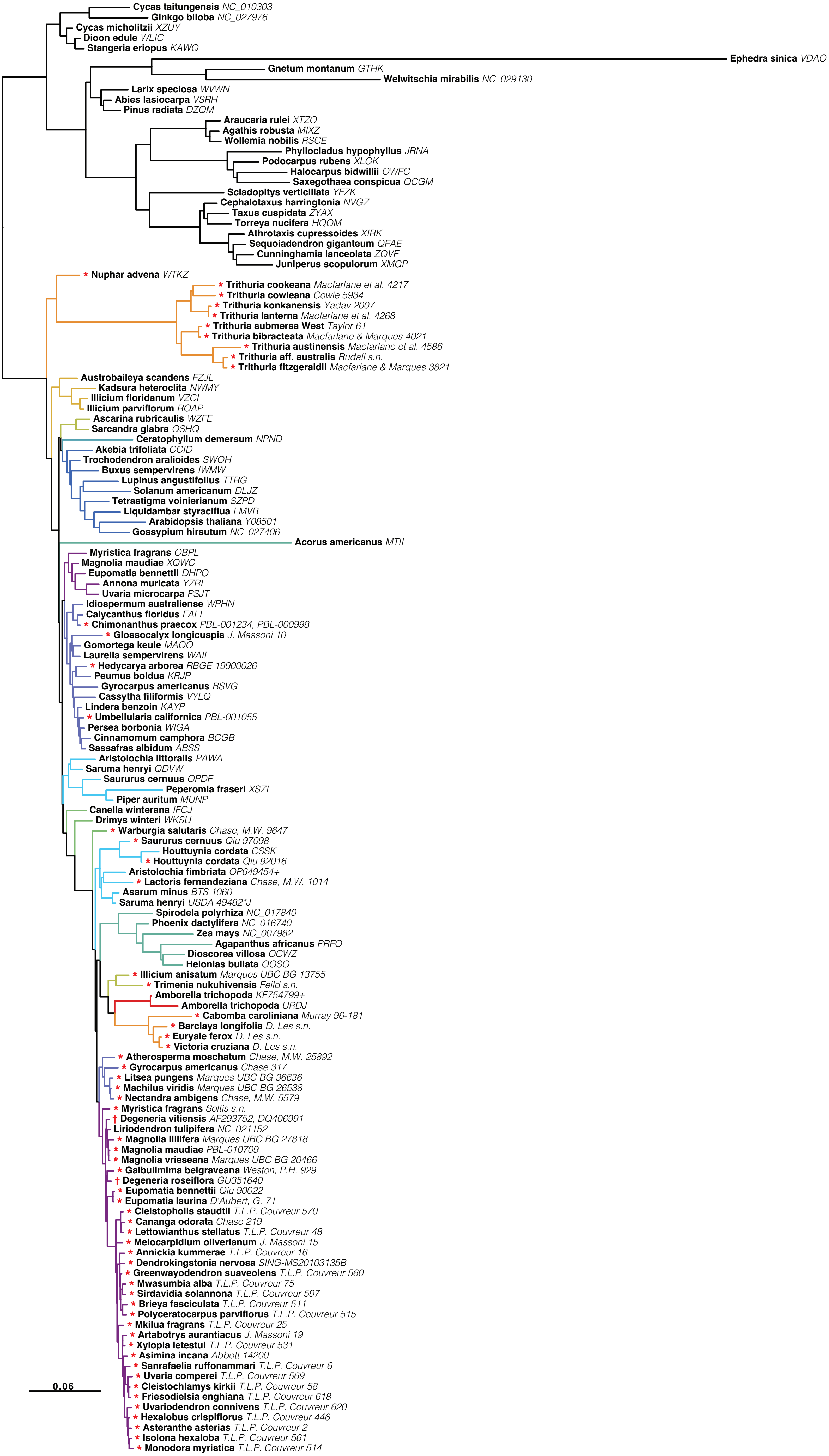

### Fig. S6

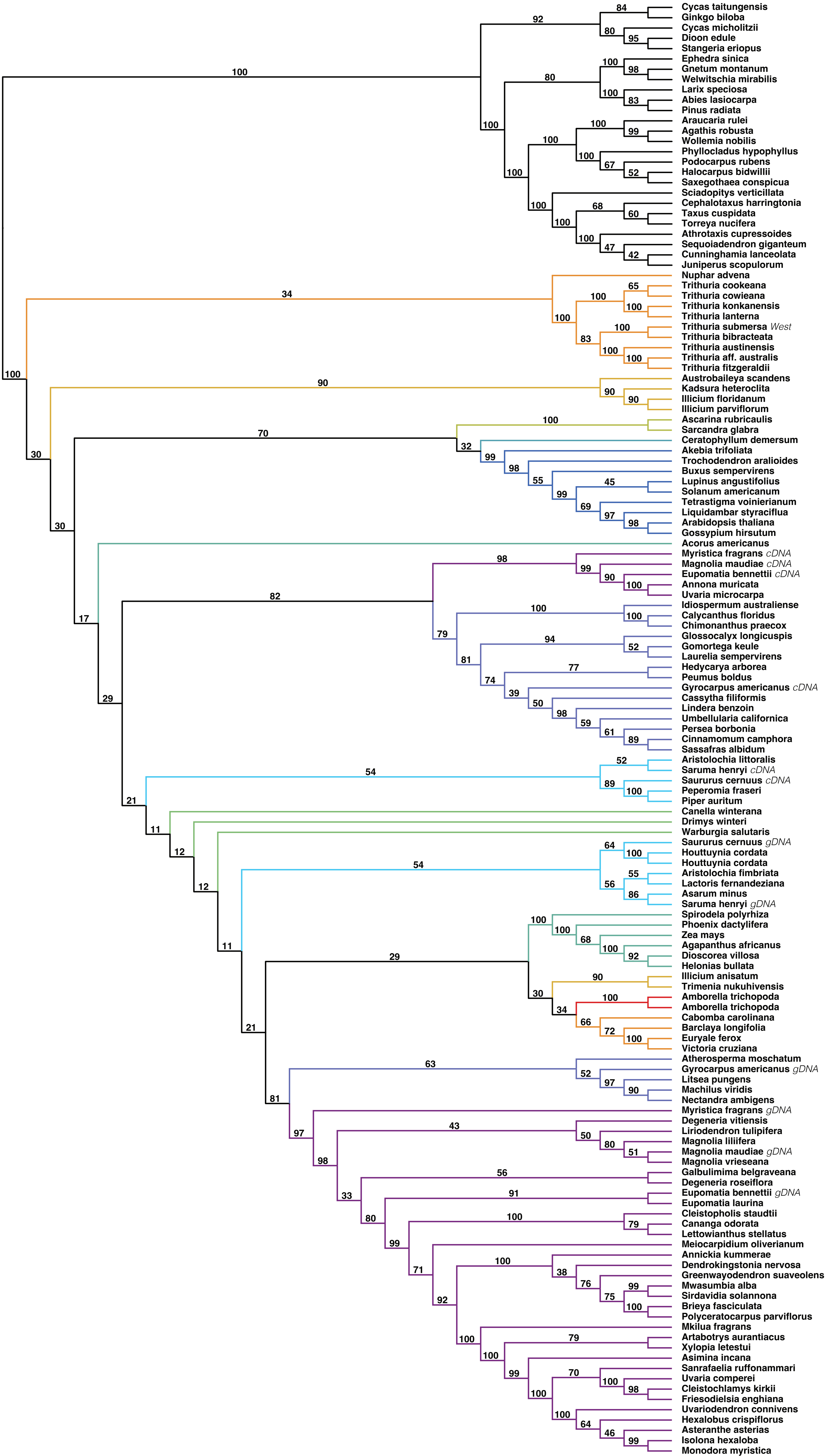

### Fig. S7

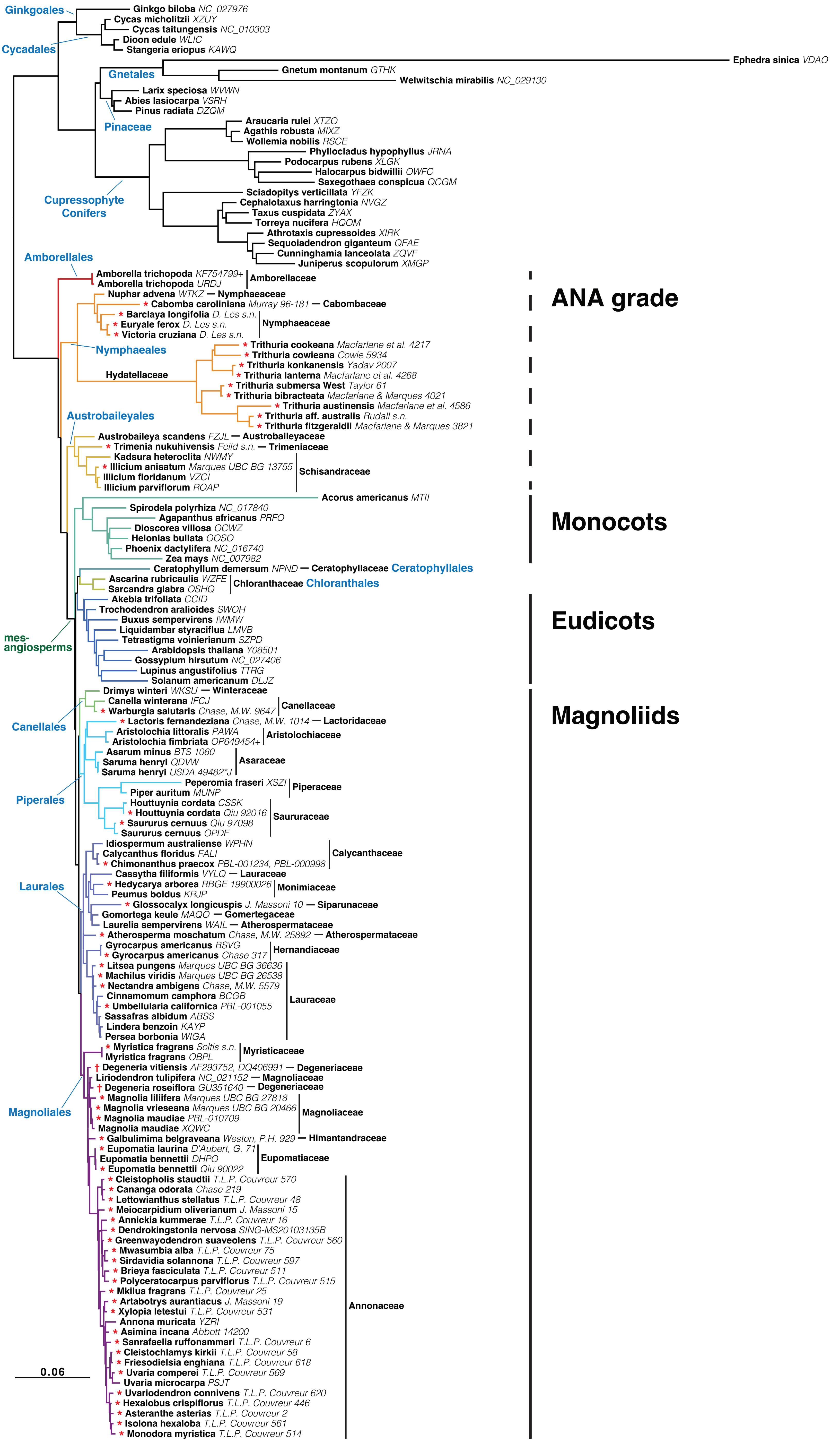

### Fig. S8

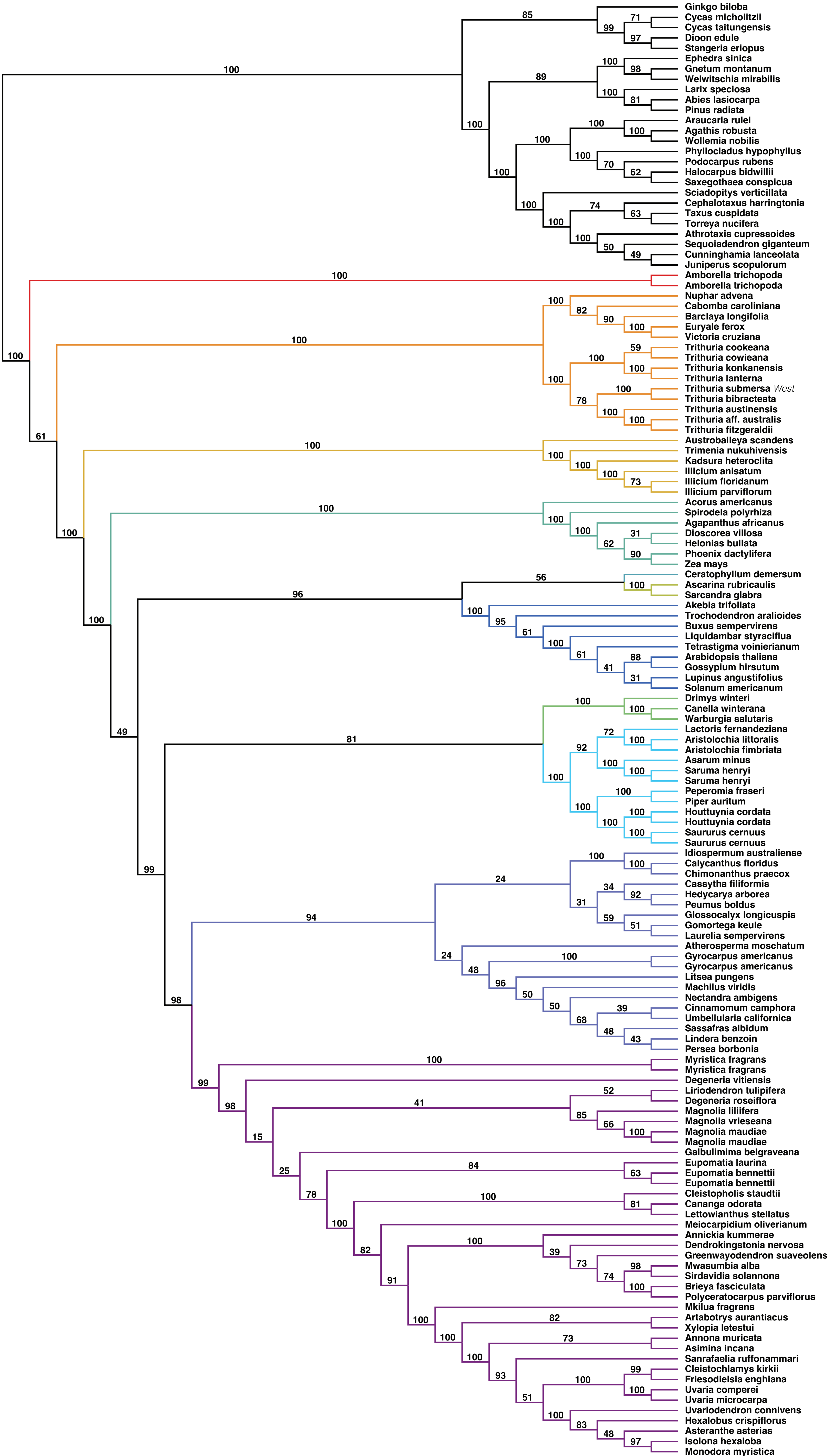

### Fig. S9

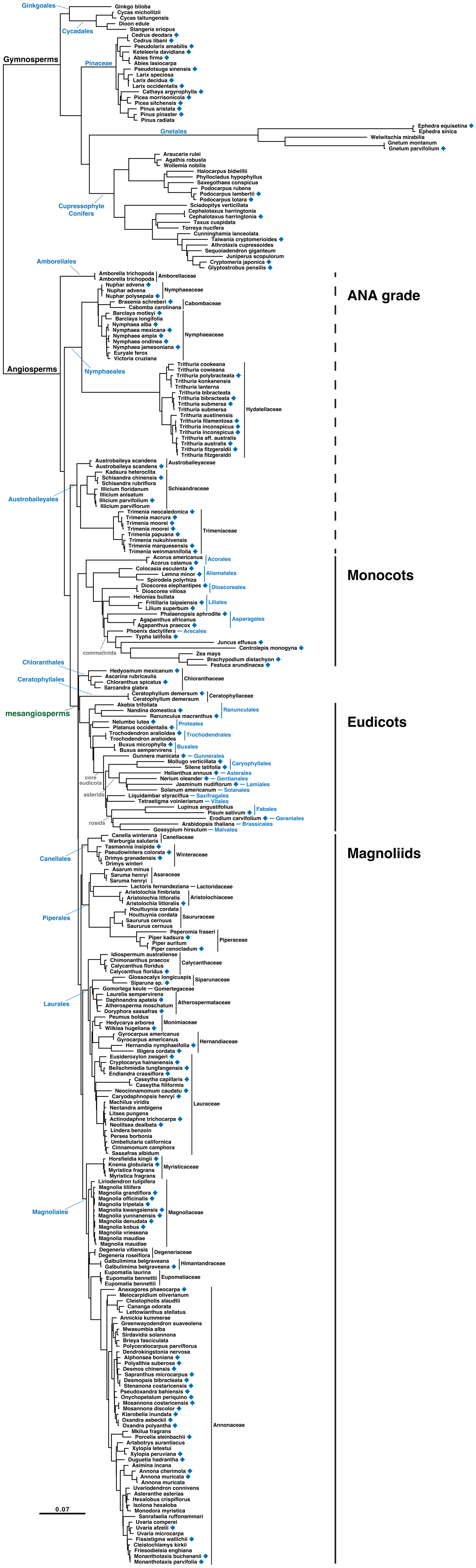

### Fig. S10

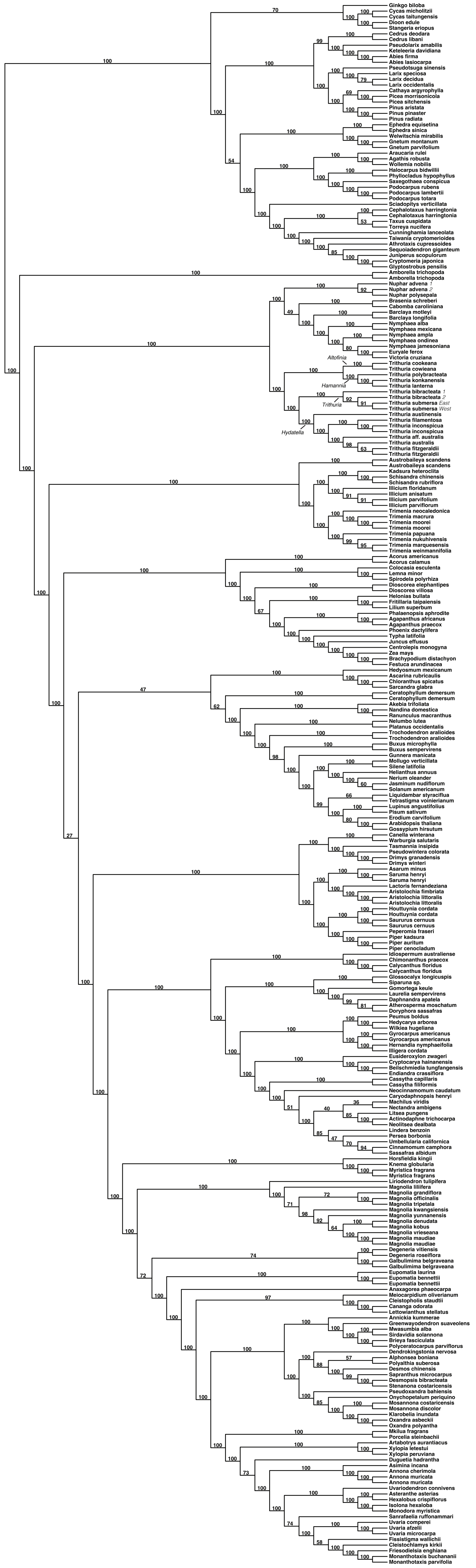
